# Molecular Display of the Animal Meta-Venome for Discovery of Novel Therapeutic Peptides

**DOI:** 10.1101/2024.05.27.595990

**Authors:** Meng-Hsuan Hsiao, Yang Miao, Zixing Liu, Konstantin Schütze, Nathachit Limjunyawong, Daphne Chun-Che Chien, Wayne Denis Monteiro, Lee-Shin Chu, William Morgenlander, Sahana Jayaraman, Sung-eun Jang, Jeffrey J. Gray, Heng Zhu, Xinzhong Dong, Martin Steinegger, H. Benjamin Larman

## Abstract

Animal venoms, distinguished by their unique structural features and potent bioactivities, represent a vast and relatively untapped reservoir of therapeutic molecules. However, limitations associated with extracting or expressing large numbers of individual venoms and venom-like molecules have precluded their therapeutic evaluation via high throughput screening. Here, we developed an innovative computational approach to design a highly diverse library of animal venoms and “metavenoms”. We employed programmable M13 hyperphage display to preserve critical disulfide-bonded structures for highly parallelized single-round biopanning with quantitation via high-throughput DNA sequencing. Our approach led to the discovery of Kunitz type domain containing proteins that target the human itch receptor Mas-related G protein-coupled receptor X4 (MRGPRX4), which plays a crucial role in itch perception. Deep learning-based structural homology mining identified two endogenous human homologs, tissue factor pathway inhibitor (TFPI) and serine peptidase inhibitor, Kunitz type 2 (SPINT2), which exhibit agonist-dependent potentiation of MRGPRX4. Highly multiplexed screening of animal venoms and metavenoms is therefore a promising approach to uncover new drug candidates.

## Introduction

Animal venoms have emerged as a vast reservoir of bioactive molecules with relatively untapped potential for drug discovery. These naturally occurring toxins, with their high potency, selectivity, and unique structural features, have evolved over millions of years to target a vast array of biological systems^1–4^. A key feature of many animal venoms is the presence of multiple disulfide bridges, which stably lock the polypeptide chain into a single conformation, resulting in resistance to protease digestion and high binding selectivity and affinity^4,5^. Recent, well-known successes of toxin-derived therapeutics include ziconotide^4,6^, an analgesic derived from cone snail venom, as well as exenatide^4,6^ and tirzepatide^7,8^, antidiabetic drugs derived from Gila monster venom. In addition, venom-derived drugs have been successfully developed for a variety of clinical applications, including anticoagulation, tumor painting^9^ and chemotherapy, underscoring the broad versatility of these natural compounds^6,10,11^.

Membrane proteins play vital roles in numerous cellular processes and are targeted by approximately 30-40% of FDA approved drugs^12^. One strategy for identifying putative therapeutics that target membrane proteins involves utilizing miniproteins, small proteins typically consisting of 20-100 amino acids, as scaffolds for drug development^13–15^. Compounds derived from these scaffolds, such as designed ankyrin repeat proteins (DARPins), affibodies and cysteine dense proteins (CDPs)^15,16^, can be optimized for specific therapeutic indications. Due to their compact size, stability, potency, and ease of engineering, peptides and miniproteins can provide especially attractive scaffolds for deriving novel membrane protein targeting therapeutics.

In pursuit of a high-throughput approach to discover novel drug candidates from diverse venom and venom-like scaffolds, we extracted all 21,311 full-length, active (mature) venom peptides and proteins contained in the UniProt database^17,18^, around 50% of which are below 90 amino acids long (∼10 kDa) and can thus be encoded via oligonucleotide library synthesis. To further expand the structural diversity of this venom scaffold library, we identified an additional 41,136 unique venom-related molecules via sequence homology searching of the Big Fantastic Database (BFD)^19^ and Serratus databases^20^, which together contain over 2.5 billion metagenomic DNA sequences. A large fraction of the animal venom polypeptide universe can therefore be encoded using molecular display for high throughput drug discovery campaigns. In this proof-of-concept study, we employed the M13 hyperphage display system, as it facilitates efficient polyvalent display of proteins with disulfide bonds^21–23^. Sequencing-assisted single-round screening can then be employed for rapid identification of target binding interactions.

As an initial demonstration of this platform, we focused on two critical receptors with distinct structural conformations: hEGFR and MRGPRX4. EGFR has a set of well-studied ligands and can be expressed both on the cell surface and in the form of a chimeric immunoglobulin fusion protein. MRGPRX4 is a seven-pass transmembrane G protein-coupled receptor (GPCR) that is expressed on the plasma membrane of primary sensory neurons in the dorsal root ganglion and play a critical role in itch sensation^24,25^. Our platform identified known ligands and novel modulators of these receptors, including the human venom-like CDPs, TFPI and SPINT2, which potentiate agonism of the human itch sensor MRGPRX4.

## Results

### Adaptation of programmable M13 hyperphage system to ligand discovery via sequencing-assisted selection

To discover novel interactions involving animal venom and venom-related ligands with diverse structural characteristics, we first optimized a programmable M13 polyvalent “hyper” bacteriophage display screening platform^21^. The M13 minor coat protein p3 is secreted through the *E. coli* periplasm where disulfide bond formation can occur, thus facilitating proper folding of fused venom-like polypeptides.

The discovery process consists of three main steps: phage library construction, single-round selection, and candidate identification via library sequencing-based analysis (Fig. 1A). To optimize binder selection efficiency, varying linker lengths were tested in the context of M13-displayed epidermal growth factor (EGF) binding to native human EGFR (hEGFR), when expressed on the cell surface at varying densities. The EGF-hEGFR interaction is known to be high affinity (Kd=0.1-1nM)^26^ and absolutely requires EGF’s three disulfide bonds. A range of linker lengths between 13 to 291 amino acids (Table S1, Fig. S1A) were tested in conjunction with three cell lines of varying EGFR expression levels: high expression (MDA-MB-468), medium expression (MDA-MB-231), and low expression (NCI-H358). As a negative control, we used CHO K1 cells overexpressing programmed cell death ligand 1 (PDL1), which are known to have low levels of EGFR expression on their surface (Fig. S1B).

**Figure 1.**
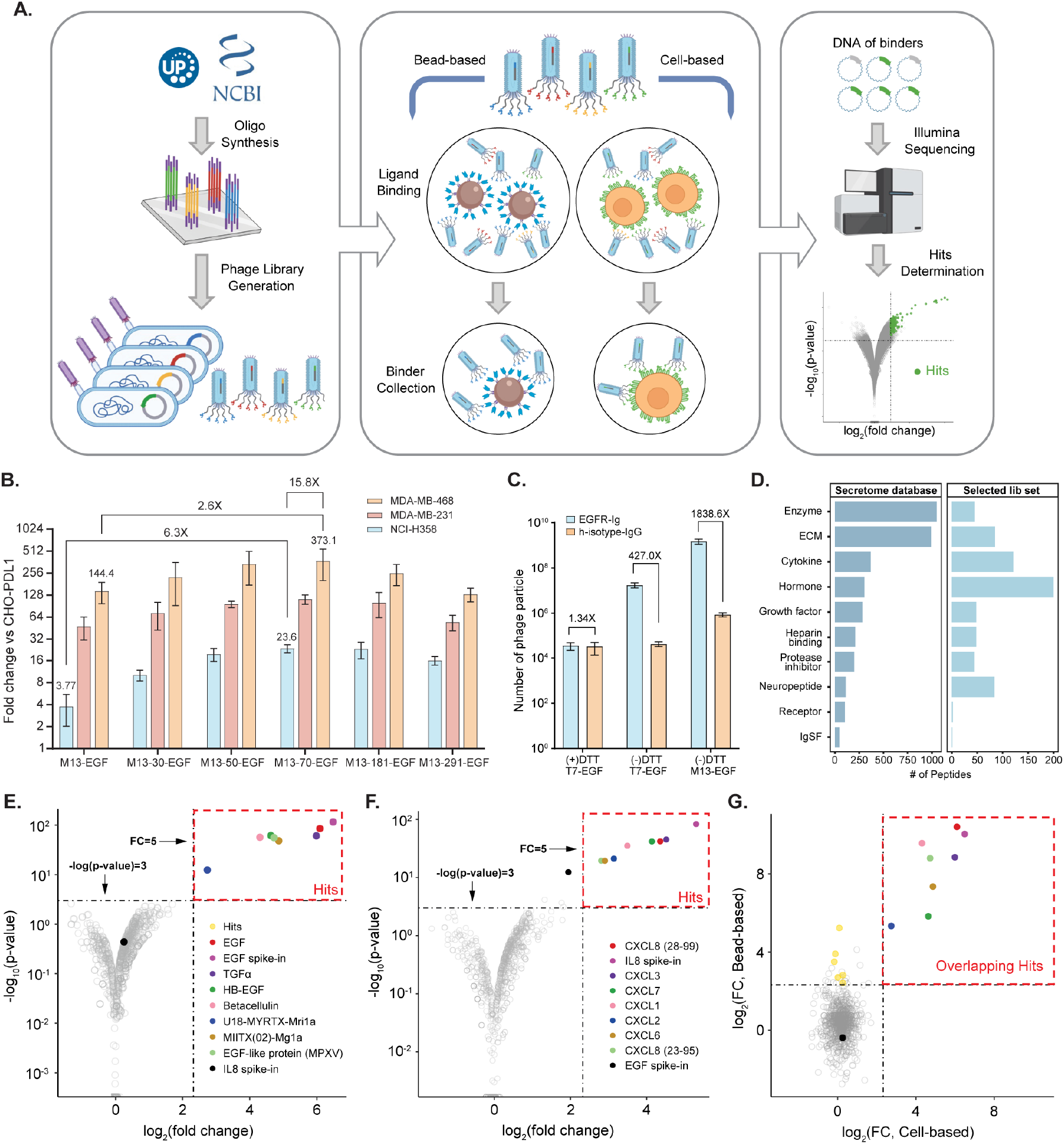
M13 polyvalent phage display workflow identifies ligands to EGFR and CXCR2. (**A**) A synthetic oligonucleotide library encoding full-length polypeptides up to 90 aa is designed and synthesized. The oligonucleotide library is cloned into the M13 hyperphage display system and grown up in *E. coli* cells (left). Proteins of interest fused with the M13 P3 protein are displayed in 5 copies on the phage surface. The phage library is screened against a target (bead based or cell based) for affinity enrichment (middle). The DNA of the enriched phage library is collected and amplified by PCR for high throughput sequencing. Informatic analysis identifies candidate binder peptides (right). (**B**) The effect of linker length and cell surface receptor expression level on M13 phage displayed EGF enrichment. (**C**) Evaluation of disulfide bond formation and its impact on EGF binding to EGFR. (**D**) The composition of the human secretome library by molecular function. ECM, extracellular matrix. IgSF, immunoglobulin superfamily. (**E-F**) Volcano plot depicting the enrichment results of M13 secretome library screening against cells overexpressing EGFR (E) or CXCR2 (F). Each point is a unique secreted peptide. Known ligands are highlighted with distinct colors. The spiked-in control is colored black, and peptides not significantly enriched are colored grey. (**G**) Scatter plot comparing peptide enrichment from EGFR-overexpressing cells (X-axis) versus Ig-EGFR coated magnetic beads (Y-axis). Each point is a unique secreted peptide from the library. Bead-only enriched peptides are colored yellow, while overlapping peptides and the spiked-in controls are colored as in (E).

The dependence of linker length was significantly more pronounced in the context of lower EGFR expression level. With a 70 aa-long linker, higher fold changes were observed across all cell lines, with approximately 16-fold higher enrichment between low (NCI-H358) and high (MDA-MB-468) EGFR expressing cells (Fig. 1B). A similar linker length-dependent trend was found when M13-displayed interleukin-8 (IL8) was tested with cells overexpressing the IL8 G-coupled protein receptor, C-X-C Motif Chemokine Receptor 2 (CXCR2) (Fig. S1C, S1D). Subsequent libraries were therefore constructed in a phagemid vector incorporating the 70 aa linker (M13-70).

The importance of the EGF disulfide bond formation was next evaluated by comparison of the M13 system with the T7 phage display system (Fig. 1C) in a bead-based Ig-EGFR binding assay (Fig. 1A). T7 phage are lytic and do not pass through the oxidizing periplasm during replication, thus forming disulfide bonds much less efficiently. EGF expressed on M13 demonstrated greatly increased binding relative to EGF expressed on T7. The lower level of T7-EGF binding was ablated by disulfide bond reduction with dithiothreitol (DTT), while T7 phage infectivity was unaffected.

### Re-discovery of known membrane receptor ligands with programmable M13 polyvalent phage display of the human secretome

To further assess how well the optimized M13 polyvalent display system could identify diverse ligands for cell surface expressed receptors, we constructed a library encompassing all 880 full-length human extracellular and secreted proteins whose mature polypeptide chains are equal to or less than 90 amino acids in length (the human secretome library). This library comprises approximately 66% of hormones listed in the Secretome database, including known EGFR ligands such as EGF, betacellulin (BTC), transforming growth factor-alpha (TGFα), and heparin-binding EGF-like growth factor (HB-EGF). We also incorporated three additional venom and venom-like proteins as positive controls for EGFR studies, namely, MPXV, an EGF-like protein from the monkeypox virus, and two ant venom toxins, Mri1a and Mg1a, which are known EGFR agonists^27^. In addition to hormones, nearly 33% of cytokines and 75% of neuropeptides from the comprehensive Secretome database are included in the human M13 secretome library. Most extracellular matrix (ECM) proteins and secreted enzymes are not included in the library due to their sizes (Table S2, Fig. 1D). Despite this size limitation, the human M13 secretome library covers a diverse range of molecular functions, including hormones (200), enzymes (45), growth factors (48), protease inhibitors (44), and neuropeptides (84) (Fig. S1E).

We employed both bead-based and cell-based panning approaches by incubating phage libraries with either magnetic beads coated with receptor extracellular domain-immunoglobulin (ECD-Ig) fusion proteins or with cells overexpressing target receptors (Fig. 1A). Screens were performed in triplicate to assess reproducibility. Candidate ligands were identified by sequencing the recovered binders after washing away unbound phage particles (Fig. 1A). Many known endogenous EGFR ligands, including EGF, BTC, HB-EGF, TGFα, Mri1a, and Mg1a, were robustly identified using HEK293T cells overexpressing EGFR (Fig. 1E). Similarly, six of the seven known endogenous ligands of the seven-pass transmembrane G protein-coupled receptor CXCR2 (CXCL1, 2, 3, 5, 6, 7, 8) were successfully detected by screening against CXCR2-overexpressing HEK293T cells (Fig. 1F). When panning against Ig-EGFR bound to magnetic beads, we identified a highly overlapping set of known and candidate ligands compared to cell-based screening, but with relatively increased fold-change values (Fig. 1G). These results establish good agreement between cell-based and bead-based screening methodologies, with enhanced signal strength and convenience for targets that can be presented at higher density on capture surfaces.

### Construction of a comprehensive animal venom scaffold library

We next designed a diverse library of all known animal protein venoms, which encompasses highly divergent target classes and bioactivities. A total of 21,311 animal venom (AV) sequences were retrieved from the UniProt database to serve as a parental set of sequences. By parsing annotations describing post-translational modifications (PTMs) and/or processing events from UniProt, we were able to extract 11,128 mature and active forms of these sequences. The remaining 10,183 sequences lacked such features. The final AV library included 10,597 mature and active sequences that were less than or equal to 90 amino acids in length (49.7% of all parental sequences, Fig. 2A, 2B).

**Figure 2.**
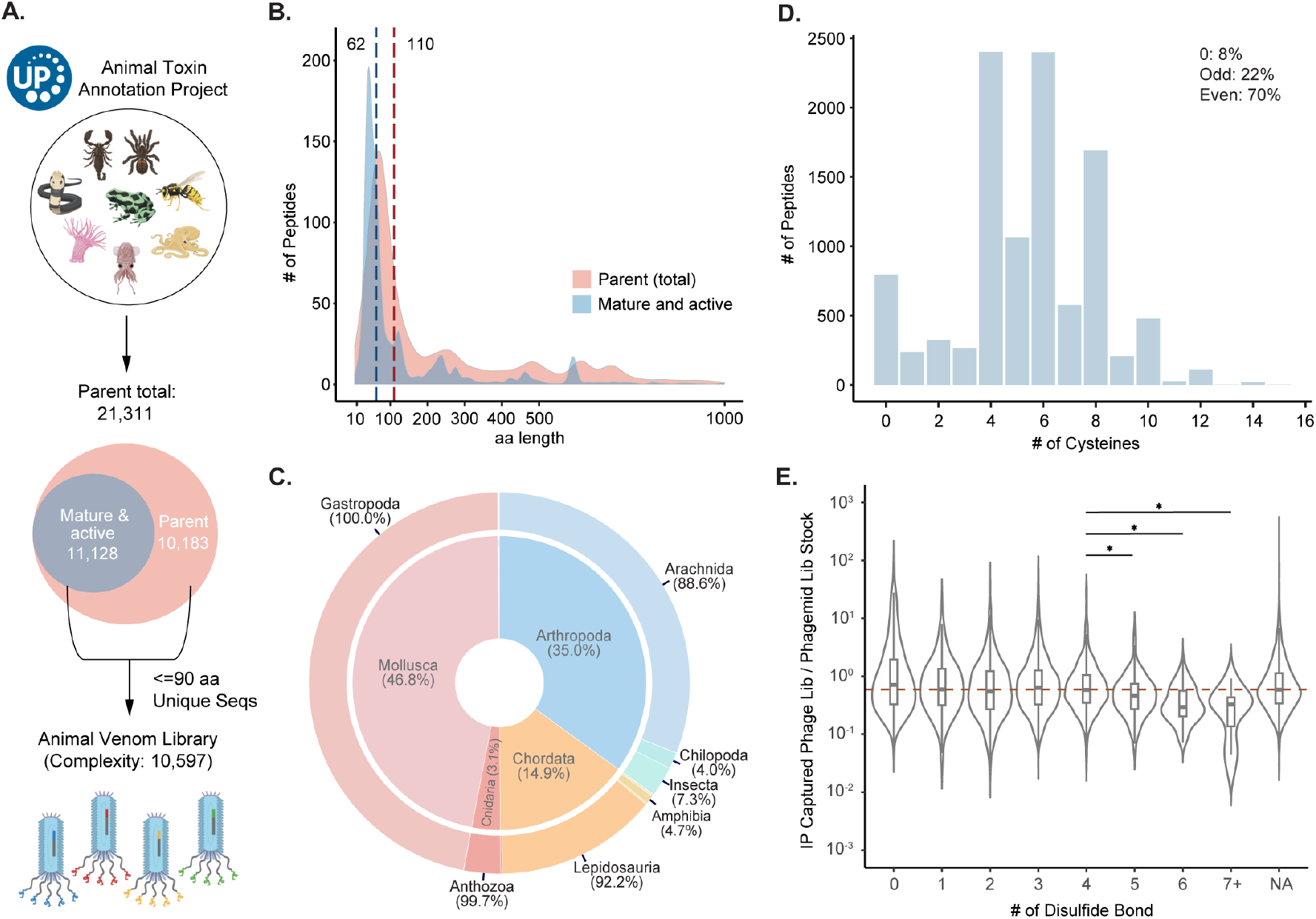
Animal venom library design and characterization. (**A**) 21,311 parent animal venom protein sequences were obtained from UniProt. Of these, 11,128 sequences possessing annotations for mature and active forms were extracted. Oligonucleotides encoding unique protein sequences up to 90 amino acids in length, regardless of their mature and active form status, comprise the M13 Animal Venom (AV) phage library. (**B**) Length distribution for AV sequences before and after the extraction of their mature and active forms. The median sequence length for the parent database is 110-aa, which reduces to 62-aa in their mature and active forms. (**C**) Composition of the AV library. The inner layer represents the animal phylum while the outer layer depicts the animal class. Table S2 provides more detailed information. (**D**) Distribution of the number of cysteines in the AV library. (**E**) Relative abundance of the AV phage library members versus the phagemid library stock as a function of disulfide bonds described in UniProt. ‘NA’ indicates sequences that possess cysteines but have unknown disulfide bond patterns. The red dashed line marks the median ratio (0.59) for the entire phage library. A two-tailed, unpaired T-test with Welch’s correction was used to assess the statistical significance of the decrease in ratio observed for library members with four or more disulfide bonds. A p-value of less than 0.05 was considered significant and is denoted by a single asterisk.

The organisms represented in the final AV library span 22 taxonomic classes, with the majority of sequences originating from cone snails (Gastropoda), scorpions and spiders (Arachnida), and snakes (Lepidosauria) (Fig. 2C, Table S3). The library displayed a broad distribution of cysteine numbers: sequences devoid of cysteines make up 8% of the total, those with an odd number of cysteines account for 22%, while a substantial majority, 70%, contain an even number of cysteines. Remarkably, 90.2% sequences contain at least two cysteines, suggesting that disulfide bond formation is likely to be a key feature of this unique and diverse scaffold library (Fig. 2D).

In the cloned phagemid library, 99.6% of the designed animal venom sequences were detectable, with 98.2% of the sequences present within one log (plus or minus) of the median. After phage packaging, 90.8% of the sequences were observed in the final phage library (Fig. S2A, S2B, Table S4), in accordance with the typical skewing and dropout of M13 phage display libraries. Factors contributing to skewness include uneven amplification, propagation and secretion of phage library members. We observed no strong correlation between the yield of specific library members containing up to four disulfide bonds, above which a slight decrease in yield was associated with increasing cysteine residues (Fig. 2E). It is important to note that the number of disulfide bonds formed during expression of the fusion proteins produced in *E. coli* may differ from the annotation of the native polypeptide.

### Expansion of the animal venom library via sequence homology searching in large metagenomic databases

To expand the diversity of our animal venom-related molecular scaffolds, we used the sequences of the animal venom library as queries to search for related sequence in the BFD and Serratus database using MMseqs2^28^ (Fig. 3A). We then removed any target sequences that were identical or longer than 100 amino acids resulting in 381,128 metagenomic sequences with venom-like attributes; referred to as the “metavenome” (Fig. 3A). To create a compact yet diverse library, we clustered the metavenome sequences at 50% sequence identity and 95% sequence length overlap using MMseqs2 and subsequently removed every sequence that overlaps with an existing animal venom sequence using the same thresholds (Fig. 3A); resulting in 41,136 representative sequences. The encoding sequences were synthesized as an oligonucleotide pool and cloned to create the M13 “metavenome” (MV) library (Fig. 3A).

**Figure 3.**
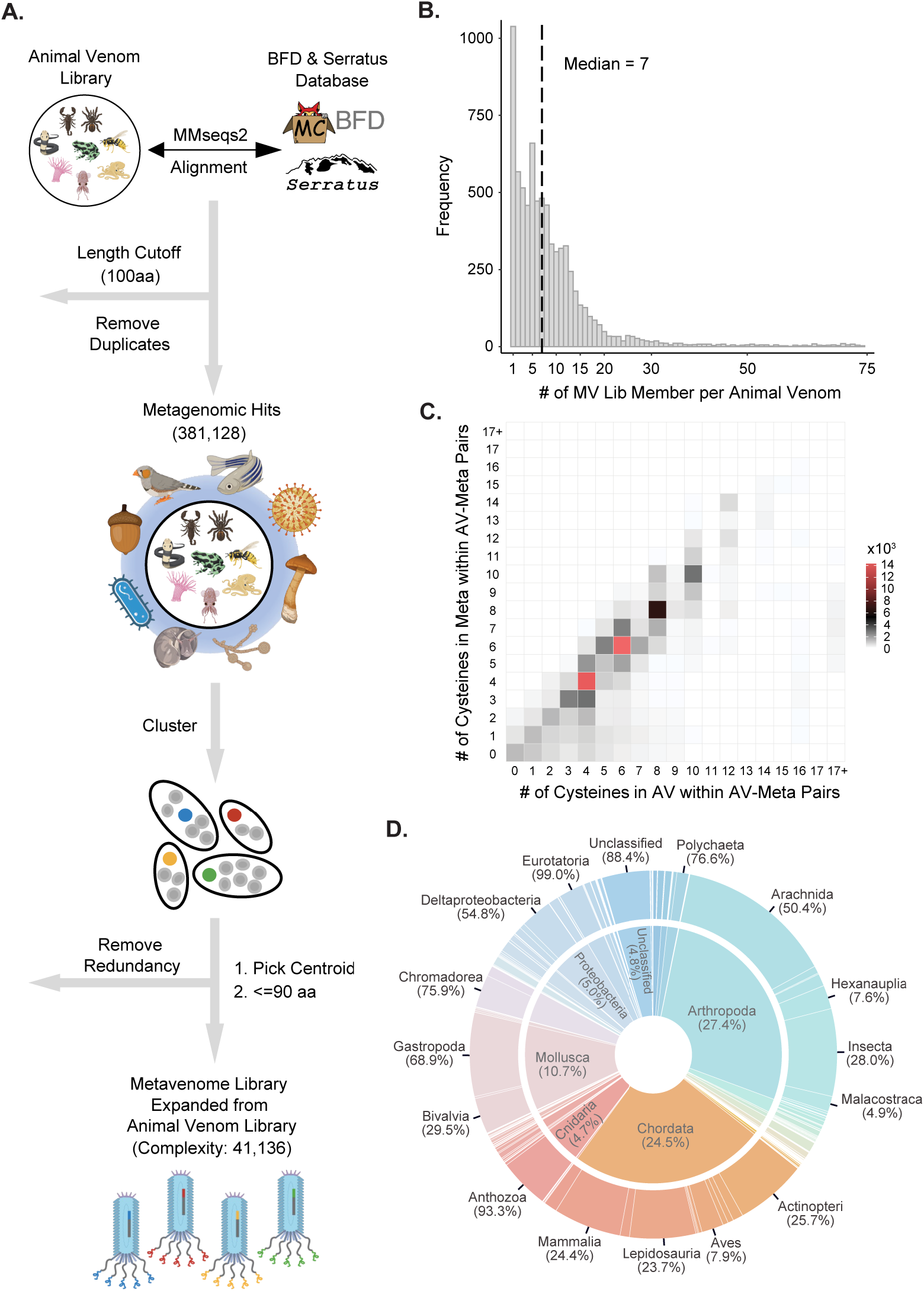
Metavenome library design and characterization. (**A**) Animal venoms were used as annotated queries to search for homologous sequences in the Big Fantastic Database (BFD) and Serratus. Identical target sequences and those exceeding 100 amino acids were removed. Metavenome sequences were clustered and excluded if they showed at least 50% sequence identity and a 95% overlap with an animal venom sequence. The final metavenome library consisted of sequences of 90 amino acids or shorter, resulting in 41,136 unique sequences that were synthesized as the metavenome library DNA pool. These sequences were subsequently cloned and packaged to form the metavenome phage library. (**B**) The number of metavenome sequences that corresponds to each animal venom. (**C**) Heat map depicting the correlation of the number of cysteines between animal venom and metavenome sequence pairs. (**D**) Composition of the metavenome library based on animal classification. The inner layer represents the animal phylum while the outer layer depicts the animal class. See Table S6 for more detailed information.

We found a median value of 7 metagenomic hits for each animal venom sequence (Fig. 3B) and importantly, a high correlation of cysteine numbers between animal venom and metavenome pairs (Fig. 3C). Whereas most other amino acids showed little or no correlation, phenylalanine and tyrosine did exhibit significant correlation, despite being less abundant (Fig. S2C, S2D). We observed no significant correlation between the yield of the phage particles and the number of cysteines in the MV protein sequences (Fig. S2E). 99.1% of the MV library members were successfully cloned into the M13-70 phagemid vector, and approximately 96% of the library members were represented in the final phage library (Fig. S2A, S2B).

Given the vast majority of sequences in the BFD and Serratus databases are not annotated, we utilized NCBI-BLAST+ to query the non-redundant (nr) database for protein annotations, including taxonomic classifications, that could be adapted to the metavenome library members. 96.3% of MV library members acquired an annotation in this way (Fig. S2F, Table S5), with a median e-value of 1.19e-15, suggesting a high level of confidence in the annotated results (Fig. S2G-J). The taxonomic diversity of the metavenome is vastly expanded compared to the AV library, comprising 130 different phyla and 210 different classes (Fig. 3D, Table S6).

### Diverse EGFR ligands discovered from the Animal Venom and Metavenome libraries

Using EGFR as our model system, we performed a comprehensive screening of the AV, MV, and secretome libraries using Ig-EGFR as the target. Candidate hits were determined by comparing sequencing results against a negative control immunoglobulin (Isotype Ig) from triplicate screens. We identified a multitude of candidate binders from all three libraries, in addition to the endogenous ligands (EGF, BTC, TGFα and HB-EGF) and previously reported animal venoms (Mg1a and Mri1a) (Fig. 4A).

**Figure 4.**
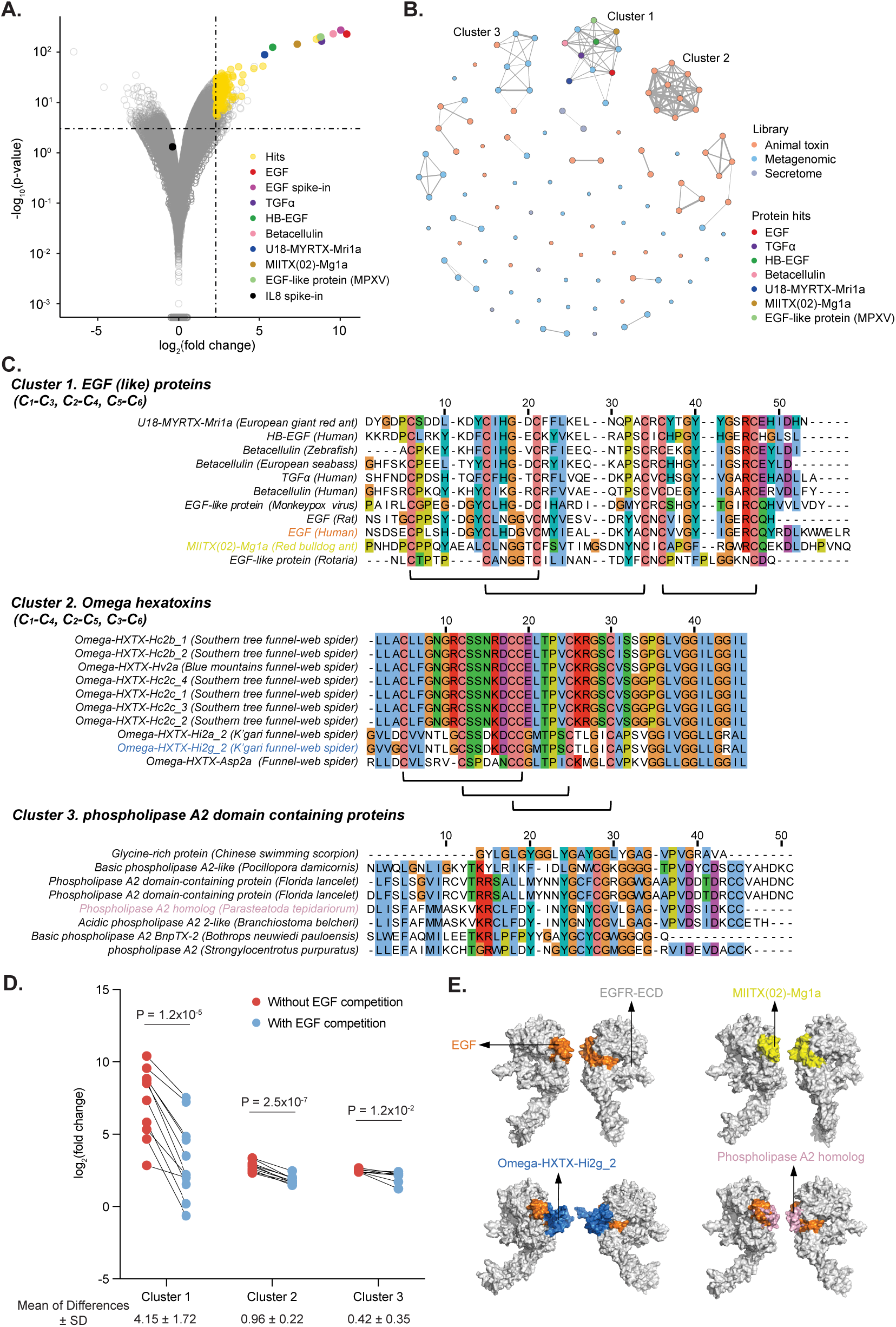
Discovery of Diverse Scaffolds and Ligands Targeting EGFR from the Secretome, AV, and MV Libraries. (**A**) Volcano plots of secretome, AV, and MV libraries screened using bead-bound EGFR-Ig. Each point is a library member, with significantly enriched clones colored yellow. Endogenous and known ligands are color-coded as in Figure 1E. (**B**) Sequence-based network graph depicting sequence homologies among candidates. Dot color corresponds to the originating library. (**C**) MSA analysis illustrates conserved amino acids shared among sequences within their respective clusters. Black underlines indicate the disulfide bond patterns provided in the UniProt database. (**D**) The fold-change values represent ligands binding to EGFR in the presence and absence of EGF in clusters 1-3. The statistical significance of the fold-change values with and without EGF was assessed using two-tailed paired Student’s t-tests. The mean differences of the fold-change values within the paired dots are displayed beneath each cluster. (**E**) Binding modes of selected ligands in each cluster predicted using RosettaDock.

We employed a sequence-based network graph approach to represent the sequence similarity among all candidate EGFR ligands. In this approach, candidate EGFR ligands, represented as nodes, were linked if they shared sequence similarity as determined by protein sequence alignment. Although many clones demonstrated no sequence homology to other hits, three dominant clusters emerged (Fig. 4B). To identify regions of sequence similarity that could underlie a shared binding mode, Clustal Omega was used to generate multiple sequence alignments (MSAs) (Fig. 4C). Among all members of cluster 1, which includes human EGF, we observed absolute conservation of the cysteine amino acids known to be crucial for EGF binding. Cluster 2 was composed exclusively of sequences from spider omega hexatoxins of the animal venom library. Cluster 2 also exhibited absolute conservation of cysteine residues, but with a disulfide bond configuration distinct from EGF. Sequences in Cluster 3 did not appear to possess disulfide bonds, and instead appeared to feature a shared phospholipase A2 domain.

Conserved sequence motifs among candidate hits suggest a shared binding mode. Clusters 2 and 3 exhibited significantly lower binding strength to EGFR versus Cluster 1. In the presence of competing EGF, the binding strengths of ligands in Cluster 1 were greatly diminished, indicative of the expected competitive binding to the receptor (Fig. S3, Fig. 4D). In contrast, EGF competition only minimally impacted binding of proteins in Clusters 2 and 3, suggestive of a distinct binding mode (Fig. 4D). To investigate this possibility, a representative from each cluster underwent RosettaDock modeling with EGFR (Fig. 4E). Ant venom (MIITX(02)-Mg1a), a known activator of EGFR signaling, was selected from Cluster 1 and accurately docked into the ligand-receptor binding pocket. However, Omega-HXTX-Hi2g_2 (Cluster 2) and Phospholipase A2 homolog (Cluster 3) docked outside of, but slightly overlapped with the ligand-receptor binding pocket (Fig. 4E), consistent with their insensitivity to EGF competition. While we could not detect antagonism in cell based assays, engineered derivatives of these lower-affinity binders may yield novel EGFR antagonists with clinical utility.

### Kunitz type domain containing proteins can potentiate MRGPRX4 signaling

The human MRGPRX4 receptor plays a crucial role in the perception of itch sensation^24,25^. Recent studies have illuminated that MRGPRX4 is activated by certain bile acids, such as ursodeoxycholic acid (UDCA). This activation triggers calcium dependent neuronal excitation, culminating in the sensation of itch^24,25^. Activation of this pathway is believed to be a key feature of liver disease-associated cholestatic pruritus, marked by elevated levels of bile acids and other endogenous metabolites in the bloodstream. Modulation of MRGPRX4 activation is therefore being pursued as a promising treatment approach.

We conducted a screening campaign using our animal venom, metavenome, and secretome libraries using cells overexpressing MRGPRX4 (HEK293-MRGPRX4) and related members of the MRGPRX family (MRGPRX1, MRGPRX2, and MRGPRX3) (Fig. 5A, 5B). Six proteins from the metavenome library demonstrated selective binding to MRGPRX4 (Fig. 5B). Interestingly, these proteins each share a highly conserved motif identifiable by multiple sequence alignment (Fig. 5C). High structural similarity among these six proteins is also predicted by AlphaFold (Fig. 5C).

**Figure 5.**
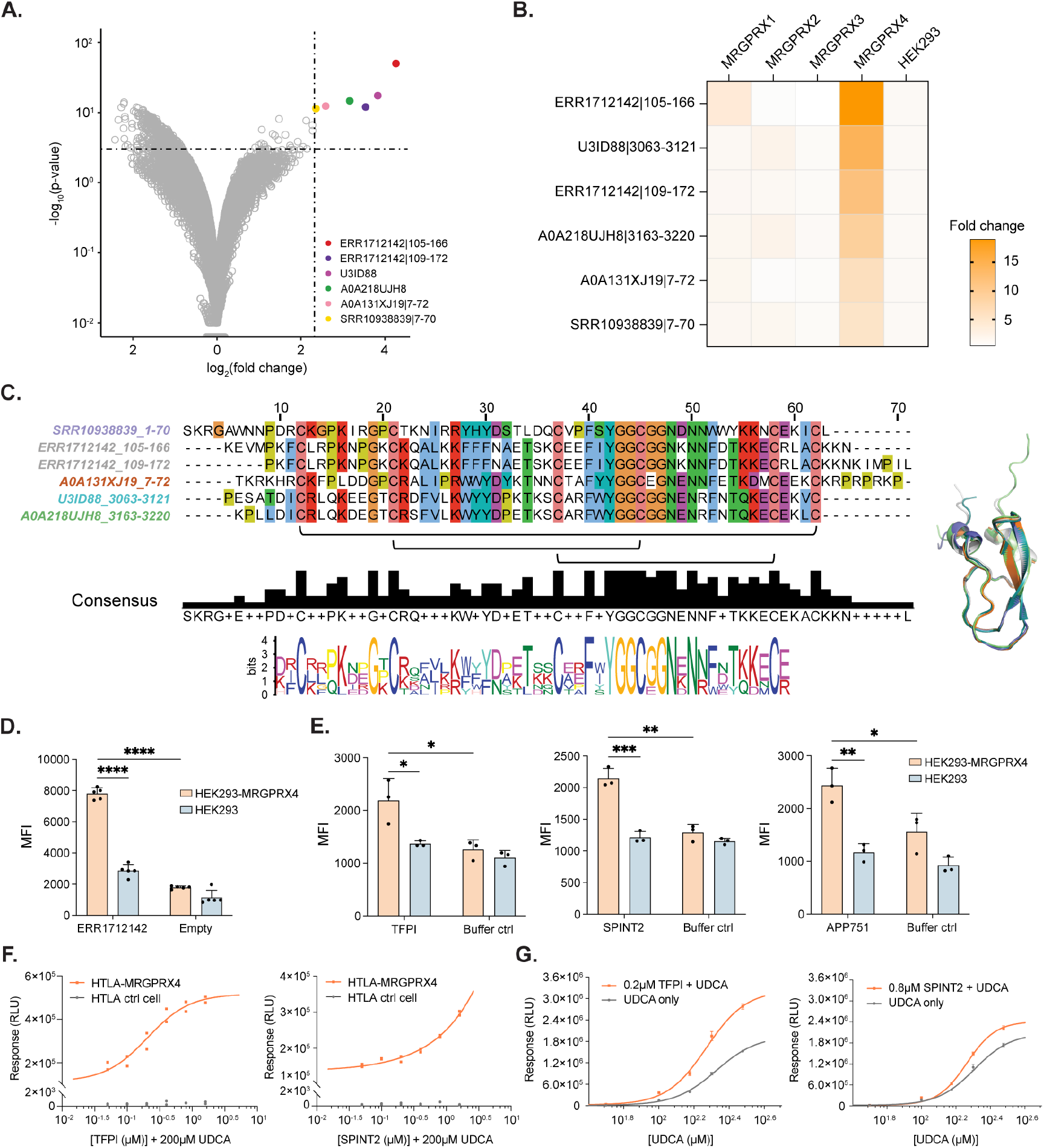
Kunitz type inhibitor is a potential potentiator for MRGPRX4 discovered from the metavenome library. (**A**) Volcano plot of animal venom, metavenome and secretome libraries binding to HEK293-MRGPRX4 cells versus HEK293 cells. Each point is a unique peptide in the AV, MV and secretome libraries. (**B**) Heat map showing selectivity of fold-change values of the 6 peptide candidates across the HEK293-MRGPRX families. (**C**) MSA analysis and motif discovery with MEME Suite identifies conserved amino acids shared among the candidate hits (left). Structural alignment of the hits using AlphaFold (right). (**D**) Flow cytometric analysis of ERR1712142 |105-166 cell binding. (**E**) Flow cytometric analysis of cell binding of TFPI (left), SPINT2 (middle), and APP751 (right). MFI, median fluorescence intensity. Data are shown as mean ± SD of 5 (in D) or 3 (in E) biological replicates. Statistical comparisons use two-tailed unpaired Student’s t-tests. ns, non-significant, *P < 0.05, **P < 0.01, ***P < 0.001, ****P < 0.0001. (**F**) Dose-response curves of TFPI (left) and SPINT2 (right) in the presence of 200 μM UDCA using the MRGPRX4 PRESTO-Tango assay. Each point is one replicate. RLU, Relative Luminescence Unit. (**G**) Dose-responses curves of UDCA in the presence of 0.2 μM TFPI (left) or 0.8 μM SPINT2 (right) in the MRGPRX4 PRESTO-Tango assay. Data are shown as mean ± SD of 3 technical replicates. RLU, Relative Luminescence Unit.

To confirm binding of putative MRGPRX4 ligands, we produced the candidate hit with the highest fold-change value – ERR1712142 |105-166 – and evaluated binding to both MRGPRX4 overexpressing cells and parent HEK293 cells. ERR1712142|105-166 exhibited significantly higher binding to the MRGPRX4 overexpressing cells compared to control HEK293 cells (Fig. 5D).

We used Foldseek^29^ to search for endogenous human homologs of these candidate MRGPRX4 binders. Foldseek interrogated AlphaFoldDB (version 4: Proteomes and Swiss-Prot), CATH clustered at 50% sequence identity, ESM Atlas-HQ, and the Protein Data Bank (PDB). All 20 of the unique human protein sequences identified as structural homologs to ERR1712142|105-166 are members of the Kunitz-type protease inhibitor class, which is characterized by six cysteines forming a distinctive disulfide bond pattern of C1-C6, C2-C4, C3-C5 (Fig. S4A, Table S7). We therefore hypothesized that Kunitz-type inhibitor domain containing proteins might modulate MRGPRX4 signaling activity.

Among the human Kunitz-type inhibitor domain-containing proteins, tissue factor pathway inhibitor (TFPI), serine peptidase inhibitor, Kunitz type 2 (SPINT2), and an isoform of amyloid-beta precursor protein (APP751) demonstrated specific binding to MRGPRX4 overexpressing cells at 30nM using a flow cytometry assay (Fig. 5E). These proteins are secreted but not included in the secretome library due to their sizes: TFPI with 254 amino acids, SPINT2 (ECD) with 169 amino acids, and APP751 with 751 amino acids. The PRESTO-Tango assay is a luciferase-based method for measuring β-arrestin recruitment to activated GPCRs, and has been previously validated for detection of MRGPRX4 activation^30^. We first conducted a cell cytotoxicity assay to ensure that prolonged exposure to these proteins would not induce cytotoxic effects (Fig. S5A). Treatment of HTLA-MRGPRX4 cells with 0.4 μM TFPI and 1.6 μM SPINT2 showed no detectable cell cytotoxicity. However, APP751 exhibited toxicity at a concentration of 0.15 μM and was therefore excluded from subsequent testing (Fig. S5A). The PRESTO-Tango assay revealed that neither TFPI nor SPINT2 activate MRGPRX4 independently (Fig. S5B). However, TFPI and SPINT2 significantly potentiate MRGPRX4 activation (Fig. S5B) in the presence of its agonist, UDCA. Dose responses of TFPI and SPINT2 were tested in the presence of UDCA at its half-maximal effective concentration (EC50) (Fig. 5F). As a potentiator, TFPI exhibited an EC_50_ of 0.2 μM, whereas SPINT2 showed a higher EC_50_ value, which was outside the dynamic range of the assay (Fig. 5F). Co-incubation of UDCA with 0.2 μM TFPI or 0.8 μM SPINT2 greatly increased MRGPRX4 activation, but did not significantly shift the EC_50_ value of UDCA, consistent with activity most likely being mediated via positive allosteric modulation (PAM) (Fig. 5G, Fig. S5C). A negative control protein, osteoprotegerin (OPG), of comparable size to TFPI and manufactured under the same conditions, showed no bioactivity towards MRGPRX4 when incubated at TFPI’s EC_50_ concentration. A second negative control protein, aprotinin, a bovine Kunitz-type domain containing protein unrelated to TFPI or SPINT2, similarly demonstrated no bioactivity in the presence of UDCA, even at its maximally achievable concentration (Fig. S5A, S5B).

## Discussion

Animal venoms and related miniproteins exhibit unique structural and functional attributes including high potency, target selectivity, and serum stability. However, the potential of these molecules has remained largely underexploited, since the capability to screen diverse venom libraries in a molecular display format has not been developed. Here we present an approach to construct and screen a comprehensive animal venom library, which was diversified to comprise the universe of sequences related to animal venoms: the “metavenome”. Single-round M13 biopanning, combined with binder detection using high throughput DNA sequencing, is shown to enable rapid discovery of lead molecular scaffolds for further optimization.

Biopanning against EGFR with the human secretome, animal venom, and metavenome libraries revealed known ligands and novel binders associated with diverse molecular scaffolds. Multiple sequence alignment of the novel EGFR binders reveals conserved residues important for binding. Separately, panning in the presence and absence of excess EGF indicates whether an interaction is likely to be competitive versus allosteric. In the case of EGFR, for example, docking simulations predict that some of the novel binders we identified have the potential to interfere with EGF signaling if further optimized.

In an effort to discover novel therapeutic scaffolds relevant to cholestatic pruritus, we undertook a comprehensive screening campaign that identified six polypeptides that selectively bound to cell surface receptor MRGPRX4, each of which appeared to contain a Kunitz-type protease inhibitor domain. AlphaFold and RosettaDock predicted their mode of binding to MRGPRX4, which involved a 21 amino acid stretch of interacting residues (Fig. S4B). In order to identify likely non-immunogenic human protein lead candidates for therapeutic development, Foldseek found several Kunitz-type protease inhibitor domain-containing structural homologs, including tissue factor pathway inhibitor (TFPI), and serine peptidase inhibitor, Kunitz type 2 (SPINT2). In cell-based functional assays, both TFPI and SPINT2 demonstrated notable potentiation of MRGPRX4 activation in the presence of its endogenous ligand UDCA. We note that MRGPRX4, TFPI, and SPINT2 do not seem to have a significant tissue co-expression pattern that might otherwise suggest a physiologic role for these interactions.

While not demonstrated here, we propose that single round, sequencing-assisted screening of venom and miniprotein libraries will be readily combined with machine learning approaches and generative artificial intelligence models to iteratively tailor library designs for specific targets or target classes^31–33^. Biopanning in the presence or absence of competitive ligands, as demonstrated here, can also be useful for guiding candidate selection and iterative library design. Advanced structural modeling and docking algorithms also provide rapidly advancing tools for scaffold engineering and lead candidate optimization.

It is important to recognize the limitations of this peptide drug discovery platform. First, the formation of disulfide bonds in *E. coli* cells may deviate from that in animal cells. This is important when considering mammalian homologs of novel binding candidates, and for downstream studies involving non-bacterial protein production systems. Second, the M13 phage display system produces libraries with significant distributional skewing. This effectively reduces the fraction of the library that is well-sampled during single-round screening campaigns. Alternative eukaryotic display systems such as yeast or mammalian display could be alternatives to the M13 phage display system. Fully *in vitro* systems like the ParalleL Analysis of Translated ORFs (PLATO)^34^ or Molecular Indexing of Proteins by Self Assembly (MIPSA)^35^ might also provide certain advantages, despite their monovalent display format. As is true for any binding-based screening technologies, molecular display cannot be used for functional or phenotypic screening.

In summary, we have developed a peptide drug discovery platform which unites polyvalent M13 phage display of cysteine-rich animal venom-derived libraries, single round sequencing-assisted selection, and deep sequence or structural modeling. We demonstrate the utility of the platform to discover ligands for both single and multi-pass membrane receptors expressed on intact human cells. Specifically, we identified known and novel EGFR binders, as well as endogenous potentiators of the itch sensor MRGPRX4. Further investigation of these molecular interactions may reveal a new strategy to block severe itch responses, such as in the setting of cholestatic pruritis. This drug discovery platform is therefore of broad utility for diverse classes of high value therapeutic targets.

## Materials and Methods

### M13 phagemid and linker design

The M13 phagemid vector was derived from the pSEX81 phagemid (PROGEN, Cat No. PR3005) modified to be compatible with next-generation sequencing (NGS) platforms (Fig.S1A). The protein of interest (POI) was cloned at the N-terminal side of the P3 protein with EcoRI and HindIII restriction sites, and a FLAG epitope was encoded downstream of the POI. To investigate the impact of linker length on ligand enrichment, a series of linkers with various lengths were incorporated between the POI and the FLAG tag. For the M13 and M13-30 constructs, a G4S linker was used directly after the POI. For the M13-50, M13-70, M13-181, and M13-291 constructs, combinations of PAS^36–38^ and G4S linkers were used to create various linker lengths between the POI and the FLAG tag. The specific linker compositions used in each construct can be found in Table S1.

### Animal venom, metavenome, and human secretome library design and synthesis

To obtain human secreted protein sequences, we accessed the UniProt database and downloaded entries based on the following search criteria: “taxonomy: ‘Homo sapiens (Human) [9606]’ (goa:(‘extracellular space [0005615]’) OR goa:(‘extracellular region [5576]’) OR locations:(location:’Secreted [SL-0243]’))”. Subsequently, we extracted mature and active sequences with annotations or labels containing “chain” or “peptide” in the “PTM/Processing” section. For DNA synthesis, we retained only sequences that were equal to or less than 90 amino acids in length, regardless of whether they have mature and active forms. In total, 880 sequences were identified, and these were reverse translated with the pepsyn library design software. Sequences less than 270 base pairs were filled with PAS linker such that the final DNA length was brought up to 300 bases when appending primer binding sequences GGAATTCCGCTGCGT and CCGAGCATTGGCACC to the 5’ and 3’ end, respectively. This systematic approach for extracting mature and active sequences from the UniProt database was also applied in the design and generation of an animal venom library.

The animal venom library, comprising full-length active (mature) animal venom and poison protein sequences, was obtained from the UniProt animal toxin database using the search terms: taxonomy:”Metazoa [33208]” (keyword:toxin OR annotation:(type:”tissue specificity” venom)). To retrieve mature and active sequences, we employed the same method as described for the human secretome library generation. We extracted sequences both with and without mature and active forms and reverse translated 10,597 unique proteins that were equal to or shorter than 90 amino acids into their corresponding DNA using the pepsyn library design software. Sequences shorter than 270 bp were supplemented with PAS linkers, and primer binding sequences GGAATTCCGCTGCGT and GTCGTGCCAGGGAAC were appended to the 5’ and 3’ ends, respectively, to achieve a final DNA length of 300 bases.

To generate a metagenomic library expanding from the animal venom sequences, we first retrieved a list of known animal venoms. We searched the UniProt database for entries containing the keywords “toxin” and “animal”. These toxins were used as queries for our searches against metagenomic databases. We searched for homologous sequences in two databases: the Big Fantastic Database (BFD) and a subset of SRA experiments identified by Serratus containing many RdRp’s assembled by Plass^39^. For the search, we used MMseqs2’s iterative search with the following parameters: 3 iterations, high sensitivity, and a maximum of 1000 hits per sequence (--num-iterations 3 -a -s 7.5 --threads 100 --max-seqs 1000 --split-memory-limit 1T). The combined number of hits from both searches was 18,184,892. We then filtered these hits to include only proteins that cover the cleaved regions of the venome to at least 90%, reducing the set to 11,120,658. Using the alignment information and cleave annotations of the query, we inferred the cleaved regions of metagenomic sequences. This process yielded 4,424,507 cleaved sequences, with 1,283,667 being unique, across 9,388 queries. Next, we removed any identical (predicted) cleaved target sequences to reduce redundancy, resulting in 2,924,870 sequences, of which 739,724 are covering 8,202 queries. We also removed predicted cleaved regions longer than 100 amino acids, due to the size restriction of the phage display method, leaving 1,380,546 pairs, with 381,128 being unique, from 6,913 queries. Next we performed two steps to reduce the sequence set size while preserving diversity of our library: (1) we removed any metagenomic sequences with at least 50% sequence identity and a 95% overlap to a cleaved venom sequence by using MMseqs2’s search; resulting in 333,787 unique pairs from 6,471 annotated queries. (2) We clustered the remaining cleaved sequences using MMseqs2 cluster^40^ with a 95% overlap and 50% sequence identity (-c 0.95 --min-seq-id 0.5), which left 70,415 pairs from 5,424 queries. To counterbalance the venoms that were only paired with a few metagenomic sequences, we added back metagenomic sequences that were removed in the last step so that each venom had at least 5 metagenomic matches, resulting in the final set of 85,406 pairs from 5,941 annotated queries and 39,583 metagenomic hits. From the metagenomic hits, we further extracted 36,140 sequences that were equal to or shorter than 90 amino acids in length. To enhance sequence diversity, we included an additional 4,996 sequences without mature and active forms and 1,000 random sequences that passed the same filtering process described above. In total, 41,136 sequences were reverse translated and supplemented with PAS linkers. To achieve a final DNA length of 300 bases, consistent with the human secretome and animal venom libraries, primer binding sequences GGAATTCCGCTGCGT and GCCTGGAGACGCCAC were appended to the 5’ and 3’ ends, respectively. The entire pipeline is implemented in python, which can be accessed at https://github.com/steineggerlab/phagedisplay-venoms. The code after the BFD/Serratus search stage runs in about 15min.

The sequences encoding the human secretome library, animal venom library, and the metagenomic library were all synthesized by Twist Bioscience (San Francisco, CA) and subsequently cloned into the M13-70 phagemid vector with EcoRI and HindIII restriction sites.

### Library cloning

M13-70 phagemid, serving as the vector for library cloning, was digested overnight with EcoRI and HindIII restriction enzymes (NEB Cat No. R3101, R3104). In order to dephosphorylate the ends, the digested vector was treated with phosphatase for a brief 10-minute duration followed by gel purification with a 2% low melting temperature agarose (ThermoFisher Cat No. 16500100) in 1X TAE buffer.

The oligo pool, comprising all three libraries used in this study, was reconstituted in molecular biology-grade water to a concentration of 100 ng/μL. A total of 10 ng of the reconstituted oligos was utilized for PCR amplification (Agilent Cat No. 600679). Two PCR reactions were carried out. In the first round of PCR, the desired library was selectively amplified by performing 10 cycles using primers with specific binding sequences unique to each library. The amplified PCR product was then column purified (QIAGEN Cat No. 28104). Following this, 10 second-round PCR reactions were set up, with each reaction containing 30 ng of purified first-round PCR product. A total of 5 cycles were conducted in the second-round PCR to introduce adaptors required for the subsequent restriction digest. Finally, the PCR-amplified product was purified again and digested with EcoRI and HindIII restriction enzymes followed by gel purification.

An adequate number of ligation reactions were prepared, each containing 50 ng of DNA (comprising both vector and insert at a 1:3 molar ratio) and high-concentration T4 DNA ligase (NEB Cat No. M0202T). The ligation mix was incubated at 16°C overnight and column-purified with molecular biology-grade water. To identify the inserts and evaluate the quality of cloning, 20 colonies from each library were picked individually and miniprepped for Sanger sequencing. Subsequent analyses showed that, on average, 50-60% of the clones were correct at the DNA sequence level, while 60-70% were accurate at the amino acid representation.

### M13 phagemid E.coli stock preparation

To ensure each library member was represented by a minimum of 100 colonies, the number of reactions for phagemid DNA library transformation into TOP10F’ electrocompetent cells (ThermoFisher Cat No. 44-0002) was calculated accordingly. The transformed cells were incubated in SOC media at 37°C for 1 hour with shaking at 250 rpm, followed by plating on LB agar plates supplemented with carbenicillin (50 mg/mL), tetracycline (5 mg/mL), and 100 mM glucose. The plates were then incubated overnight at 30°C, and the phagemid library stock were then collected and stored in -80°C with LB supplemented with 25% glycerol, carbenicillin (50 mg/mL), and 100 mM glucose. To generate a single clone phagemid construct, the same transformation and plating protocols were followed as for the library.

### M13 phage expression, purification, and quantification

For phage library production, the *E. coli* phagemid library stock, which covered at least 100-fold of its library complexity, was diluted in pre-warmed LB media supplemented with 100 μg/mL carbenicillin and 100 mM glucose to an OD600 of 0.1. The culture was then incubated at 30°C until it reached an OD600 of 0.4. Upon reaching the desired OD, the cells were infected with hyperphage (PROGEN Cat No. PRHYPE) at a multiplicity of infection (MOI) of 20 and incubated at room temperature for 15-20 minutes without shaking. The cells were subsequently incubated at 30°C with shaking at 250 rpm for 45 minutes. The hyperphage-infected cell pellet was collected by centrifugation at 4,000 g and resuspended in 2XYT media supplemented with 100 μg/ml carbenicillin, 50 μg/ml kanamycin, and 200 μM IPTG to induce protein production. To facilitate phage library production, the cell culture was incubated for 16 hours at 30°C with shaking at 230 rpm. The bacteria were then pelleted by centrifugation at 8,000 rpm and 4°C, and the phage library was collected by filtering through a protein low binding 0.22 μm filter. To remove any residual bacterial debris, the library underwent two rounds of centrifugation at 10,000 g for 5 minutes at 4°C.

Subsequently, the phage library was concentrated to a volume at least ten times smaller than its original volume and buffer exchanged with PBS using a 15 mL Amicon 50 kDa spin filter. A protease inhibitor was added to the PBS-based concentrated phage library. To determine the number of phage particles in the solution, 100 μL of the phage library was subjected to small-scale dialysis with DNAse I buffer (10mM Tris-HCl, 2.5mM MgCl_2_, 0.5mM CaCl_2_) overnight at 4°C. The following day, DNAse I was added to the dialyzed solution and incubated at 37°C for 10 minutes to destroy any single and double-stranded DNA that was not packaged within the M13 phagemid particle.

Finally, DNAse I was heat inactivated at 75°C, and the solution was diluted 100-fold with molecular biology-grade water before quantifying the number of phage particles using quantitative polymerase chain reaction (qPCR).

### Metagenomic library annotation

In this study, we aimed to annotate protein information and assign taxonomic classifications for the metagenomic library using NCBI-BLAST+ software (version 2.13.0). Peptide sequences went through a blastp search against the comprehensive NCBI non-redundant protein database (nr, as of July 2022) to find similarities in sequence. Depending on the query length, we tailored the alignment parameters accordingly. For sequences that were equal or less than 30 amino acids in length, we employed the parameters: “-evalue 200000 - max_target_seqs 10 -max_hsps 1”. The e-value was adjusted to accommodate short sequences. For amino acid sequences ranging between 31 and 90, the parameters were: “-evalue 1e-3 - word_size 6 -max_target_seqs 10 -max_hsps 1” (default parameters: “-evalue 10 -word_size 3 - max_target_seqs 500 -max_hsps >=1”). All other blastp parameters were maintained at default settings. We had explored more sensitive parameters, such as smaller word sizes, but ultimately selected the current parameters to achieve a fine balance between sensitivity and search runtime. The blastp output included alignment metrics, protein names and accession IDs, and taxonomy identifiers (TaxIds). Utilizing this output, we identified the best hits based on the lowest E-value and queried the corresponding TaxIds within Taxonkit^41^ to annotate the complete taxonomic lineages.

### Cell lines and culture

The NCI-H358 and HEK 293T-CXCR2 cell lines were generously provided from Prof. Bert Vogelstein (Department of Oncology, Pathology, and Molecular Biology and Genetics, Johns Hopkins University) and Prof. Jamie Spangler (Department of Chemical and Biomolecular Engineering, Johns Hopkins University) respectively. NCI-H358 cells were cultured in McCoy’s 5A medium supplemented with 10% heat-inactivated fetal bovine serum (FBS) and 50 U/ml penicillin-streptomycin, while MDA-MB-468, MDA-MB-231 overexpressing EGFR, and HEK 293T cells overexpressing EGFR and CXCR2 were maintained in complete DMEM culture media (high-glucose DMEM supplemented with sodium pyruvate, 10% heat-inactivated FBS, and 50 U/ml penicillin-streptomycin). HTLA cells, stably expressing a tTA-dependent luciferase reporter and a β-arrestin-TEV protease fusion gene, were maintained in complete DMEM media supplemented with 2 μg/ml Puromycin and 100 μg/ml Hygromycin. HTLA cells stably expressing MRGPRX4-tango were maintained in complete DMEM media supplemented with 2 μg/ml Puromycin, 100 μg/ml Hygromycin, and 100 μg/ml Zeocin. All cell lines were incubated at 37 °C, 95% humidity, and 5% CO2.

### Generation of an EGFR-overexpressing stable HEK 293T cell line

To generate HEK 293T cells overexpressing EGFR, a lentiviral transduction method was employed. The full-length EGFR gene was inserted into the pCDH lentiviral expression vector, and the lentiviral particles were produced using the pPACKH1 HIV Lentivector Packaging System (System Bioscience Cat No. LA500A-1). In brief, 3E+06 HEK 293T cells were plated on a 10 cm dish and incubated overnight in Iscove’s Modified Dulbecco’s Media (IMDM) containing 10% FBS and 2 mM L-glutamine. On the subsequent day, 2 μg of the pCDH plasmids encoding EGFR were co-transfected with the pPACK packaging plasmid mixture into HEK 293T cells, utilizing GeneJuice (Sigma Cat No. 70967) as the transfection agent. After 48 hours, the media containing EGFR lentivirus was collected, filtered through a 0.45 μm filter, and used for transduction of 1E+05 HEK 293T cells in a 24-well plate along with 8 μg/mL polybrene (Sigma Cat No. TR-1003-G) in 500 μL of complete DMEM. Following transduction, the cells were subjected to centrifugation at 800×g for 30 minutes at 32°C and incubated overnight at 37°C under 5% CO_2_ conditions in a humidified incubator. The next day, the culture media was replaced with fresh complete DMEM, and the transduced cells were collected 10 days post-transduction to evaluate EGFR expression using flow cytometry. Cells exhibiting the top 10% of EGFR expression were sorted, collected, and subsequently cultured in complete DMEM.

### Flow cytometry analysis of receptor expression level

Flow cytometry was utilized to evaluate the expression levels of target receptors in harvested cells. Approximately 5E+05 cells were resuspended in 1 mL FACS washing buffer (1% BSA or 2% serum in PBS without Ca^2+^ or Mg^2+^, containing 0.05% NaN_3_) and transferred to a FACs tube. Following centrifugation at 300 g for 5 minutes, the supernatant was discarded. Cells were then incubated with 1 μg of primary antibody diluted in 100 μL of FACS buffer. For EGFR and CXCR2 overexpressing cells, mouse anti-EGFR mAb (ThermoFisher Cat No. MA5-13070) and mouse anti-CXCR2 mAb (R&D Systems Cat No. MAB331) were used for primary antibody staining, respectively. Samples were then stained for at least 30 minutes on ice before washing with 2-5 mL FACS buffer. After two subsequent washes by centrifugation at 300 g for 5 minutes at room temperature, cells were incubated with goat anti-mouse Alexa Fluor 488 mAb secondary reagent (ThermoFisher Cat No. A11001) for at least 30 minutes ice, shielded from light. Following a final double wash with FACS buffer, cells were adjusted to a concentration of approximately 1E+06 cells/mL, and the expression levels of target receptors were analyzed using flow cytometry.

### Single-round cell-based and bead-based screening

Two distinct screening methods were employed to identify novel ligands: a cell-based system and a receptor-Fc chimera immunoprecipitation technique. Both approaches involved incubating phage libraries with target receptors, collecting binders, performing PCR, and determining candidate hits based on next-generation sequencing (NGS) results. To ensure statistical validation and reproducibility, each condition was represented by triplicate samples.

In the cell-based screening method, cells overexpressing target receptors were detached with cell dissociation buffer (Corning Cat No. 25-056-CI) and collected after passing through 40 μm cell strainer. The collected cells were washed three times with ice-cold PBS containing 1% (w/v) BSA (PBSA) and then incubated separately with the phage library in 1.5 mL Eppendorf tubes containing 1% PBSA for 30 minutes at 4°C to minimize non-specific binding. Approximately 1E+06 cells were subsequently incubated with the phage library in 1% PBSA for 4 hours at 4°C on an end-to-end rotator. After washing the cells three times with ice-cold 1% PBSA, ssDNA of the bound phages was collected using a modified protocol from the Quick-DNA/RNA Miniprep Kit (Zymo Cat No. D7011). Briefly, after cell lysis and chromosomal DNA removal, RNAse A (ThermoFisher Cat No. EN0531) was added to digest cellular RNA at 37°C for 30 minutes. Next, the RNAse-digested solution was mixed thoroughly with an equal volume of 40% molecular biology-grade ethanol for ssDNA phagemid capture. The standard column purification protocol from the Quick-DNA/RNA kit was then applied, and 15 μL of eluted ssDNA binders were stored at -20°C for subsequent quantification or sequencing.

For the phage immunoprecipitation-based ligand discovery, 1 μg of EGFR-Fc (R&D Systems Cat No. 344-ER) and Etanercept chimeric proteins (Millipore Sigma Cat No. 185243-69-0), representing the extracellular domain of the target receptor fused to the Fc region of immunoglobulin G, was incubated with the pre-blocked phage library overnight at 4°C. The following day, 20 μL of protein G beads (ThermoFisher Cat No. 10009D) were added to the phage-chimera mixture and rotated for 4 hours at 4°C to immunocapture (IP) all chimeras. The beads were then washed three times with PBS containing 0.01% NP-40 and stored in -80°C before sequencing.

### NGS sequencing and data analysis

Binders from both cell-based and bead-based screening methods were subjected to PCR for library insert amplification and sample-specific barcode incorporation. The PCR products were pooled and analyzed using Illumina sequencing. The protocol employed here is a standard PhIP-Seq protocol that has been described in detail^42^. Briefly, a first PCR was performed with primers that flank the displayed peptide inserts, and a subsequent PCR added adapters and sample indexes for single-end dual index Illumina sequencing.

Illumina sequencing FASTQ outputs were demultiplexed and mapped to the reference sequences through alignment. Perfect matches were counted to generate a read count matrix, with rows representing polypeptides in the library and columns corresponding to samples including the targets of interest and relevant negative controls. Each sample, including the negative control, was assayed in triplicate. For the cell-based screening method, the negative controls were parental, non-transduced cells. For the phage immunoprecipitation-based method, the negative controls were human isotype immunoglobulin G (hIgG). EdgeR software^43,44^ was adapted to calculate the maximum likelihood fold-changes and the p-values of differential abundance in order to assess the enrichment for each sample relative to the negative controls. We determined that a significantly enriched polypeptide (“hit”) should have a p-value less than 0.001, a fold-change of at least 5, and a read count of at least 15 in two of the triplicate samples (fold change values and p values were calculated with the EdgeR software). These criteria were established heuristically to achieve optimal sensitivity and reproducibility.

### Sequence-based network graph analysis and MSA evaluation

Network graph analyses were performed using R 4.2.0, with the network graphs generated through the R iGraph software package. Enriched peptides for each receptor screened were chosen to create the corresponding network graphs based on their sequence homology. Connectivity between sequences was determined by aligning protein sequences with blastp, utilizing the rBLAST package interfacing with the ncbi-blast+ (2.13.0) software. Alignment parameters included “-evalue 1e-3 -max_hsps 1 -seg no -soft_masking false -word_size 5 - max_target_seqs 100000 -comp_based_stats none”. In the network graphs, polypeptides were represented by nodes, and those sharing sequence similarity while meeting alignment thresholds (E value < 0.001) were connected. The link width was proportional to the BLAST bit-scores. To explore potential conserved motifs, sequences from individual clusters with more than two peptides were extracted for multiple sequence alignment using the Clustal Omega tool provided by the European Molecular Biology Laboratory’s European Bioinformatics Institute (EMBL-EBI). The same sequences were also employed for motif discovery using MEME Suite 5.4.1.

### Binding assay for ERR1712142 |105-166 and human structural homologs screening against MRGPRX4

pLicC-MBP-ERR1712142 plasmid was extracted from bacterial lysate with Maxiprep (Qiagen Cat No. 12662) after overnight incubation in BL21 strain BL21 (λDE3) pLysS. The plasmids were transcribed in-vitro using the HiScribe T7 High Yield RNA Synthesis Kit (New England Biolabs Cat No. E2050S) to produce RNA. The 40 μL reaction contained 500 ng plasmid template, 20 μL NTP buffer mix and 4 μL T7 RNA polymerase and was incubated at 37 °C for 2 hours. After transcription the product was diluted with 60 μL molecular biology grade water DNA and the plasmid was cleaved at 37 °C for 15 minutes by the addition of DNAse I. Then 50 μL of 1 M LiCl was added to the solution and incubated at -20°C for 30 minutes. The centrifuge was cooled to 4°C, and the RNA was spun at maximum speed for 30 minutes. The supernatant was removed, and the RNA pellet washed with 70% ethanol. The sample was spun down at 4°C for another 10 minutes, and the 70% ethanol removed. The pellet was dried at room temperature for 15 minutes, and subsequently resuspended in 100 μL water.

RNA of translated using the PURExpress ΔRibosome Kit (New England Biolabs Cat No. E3313S). The translation reactions contained 0.4 μM mRNA, 10 μL Solution A, 3 μL Factor Mix, 0.3 μM Ribosomes, 20 U Murine RNase inhibitor (Protector RNase inhibitor, Millipore Sigma Cat No. 3335399001), 1 μL of Disulfide Bond Enhancer 1 and 1 μL of Disulfide Bond Enhancer 2 (New England Biolabs Cat No. E6820S). The reactions were incubated at 37 °C for 8 hours and used immediately or stored at -80 °C. 2.5 μL of the translated product was run on a 4-12% Bis-Tris SDS-PAGE gel and transferred onto a PVDF membrane and stained for validation of translation.

HEK293-MRGPRX4 and HEK293 cell lines were detached with cell dissociation buffer and collected after passing through 40 μm cell strainer. The collected cells were washed three times with ice-cold 1% PBSA. The cells were then incubated with ERR1712142, 30nM TFPI (Acro Biosystems, Cat No. TFI-H5226), 30nM SPINT2 (Sino Biological, Cat No. 10324-H08H) and 30nM APP751 (BioLegend, Cat No. 842601) prepared in 1% PBSA for 4h at 4°C on an end-to-end rotator. After washing the cells three times with ice-cold 1% PBSA, the cells were washed one more time with FACS washing buffer and then incubated with PE or Alexa647-conjugated FLAG tag antibody (PE, BioLegend Cat No. 637309, 1:150 dilution; or Alexa-647, ThermoFisher Cat No. MA1-142-A647, 1:400 dilution) for ERR1712142 or His tag antibody (ThermoFisher Cat No. MA1-21315-A647, 1:1000 dilution) for human homologs for 10-15min at room temperature in the dark. The cells were then washed three times with ice-cold FACS washing buffer and resuspended in 400 μL buffer for the following flow cytometry analysis. Flow cytometry analysis was conducted on Beckman Coulter CytoFLEX. Data from flow cytometry was processed on Flowjo software.

### PRESTO-Tango assay

The HTLA-MRGPRX4 and HTLA cells were seeded at a density of 1.3E+04 cells/well in a 96-well half-area flat-bottom opaque white plate that was pre-treated with 0.1 mg/ml poly-D-lysine. The following day, cells were stimulated by addition of UDCA at certain concentrations. For the modulator activity test, the cells were preincubated with TFPI, SPINT2, APP751, Osteoprotegerin (Acro Biosystems, Cat No. TNB-H5220), or aprotinin (Sigma-Aldrich, Cat No. A1153) for 1 hour at 37°C before addition of UDCA. The cells were then incubated for 12 hours at 37°C. After incubation, the medium was removed, and 25 μL per well Bright-Glo reagent (Promega Corporation, Cat No. E2620) was added. The plates were incubated for 5 min at room temperature in the dark. The luminescence was measured by Flexstation 3 system (Molecular Devices). The dose-response curves were generated using Graphpad Prism 9 software.

### Lactate dehydrogenase (LDH) cytotoxicity assay

Cytotoxicity of proteins was measured by CyQUANT LDH cytotoxicity assay kit (Invitrogen, Cat No. C20300) following the manufacturer’s instructions. Briefly, the HTLA-MRGPRX4 and HTLA cells were seeded and treated by the same procedure as the PRESTO-tango assay. After treatment, 40μL medium of each well was transferred to a 96-well flat-bottom plate for reactions. Additionally, cell lysis solution was added into untreated cells as lysed cell controls, and after complete lysis, 40μL supernatant was transferred to the reaction plate. 40μL per well reaction mixture was added to the reaction plate. The plates were incubated at room temperature for 30min in the dark, and then 40μL stop solution was added into each well. The absorbance at 490nm and 680nm was measured by Varioskan LUX Multimode Microplate Reader (Thermo Scientific). LDH activity was calculated by subtracting the 680nm absorbance value from the 490nm absorbance value. For calculation of % cytotoxicity, LDH activity of the untreated wells was subtracted from LDH activity of both treated wells and lysed cell controls, then LDH activity of treated wells was divided by activity of lysed cell controls.

### Docking prediction

We employed the AlphaFold-Multimer^45^ implementation in ColabFold^46^ version 1.5.2, incorporating templates from PDB70, to perform docking of four targets on EGFR (EGF, MIITX(02)-Mg1a, omega-HXTX-Hi2g_2, and phospholipase A2 homolog) and ERR1712142 |105-166 on MRGPRX4. AlphaFold-Multimer generated five docked models for each target, of which we selected the top-ranked model, guided by the average pLDDT scores. We further subjected the top-ranked model to RosettaDock 4.0^47^, using conformational ensembles on ROSIE^48^. From the 1000 models generated by RosettaDock, we selected the top 10 based on the Rosetta interface scores. To identify key amino acids involved in binding, we generated contact maps of the top-ranked models from both AlphaFold and RosettaDock using MAPIYA, considering the distance between amino acids with a cut-off set at 5 Å^49^.

## Acknowledgments

This work was made possible by support from NIH grant GM136724 (H.B.L.). We thank J. Spangler for generously providing the HEK 293T-CXCR2 cell line. We thank B. Vogelstein for generously providing the MDA-MB-231 and NCI-H358 cell lines. We thank D. Wirtz for generously providing the MDA-MB-468 cell line. We also thank L. Orzolek and H. Hao of the Johns Hopkins Transcriptomics and Deep Sequencing Core Facility for all their help with next-generation sequencing. Parts of some figures were created with BioRender.com. M.S. acknowledges support from the National Research Foundation of Korea (grants 2019R1A6A1A10073437, 2020M3A9G7103933, 2021R1C1C102065, and 2021M3A9I4021220), the Samsung DS Research Fund, and the Creative-Pioneering Researchers Program through Seoul National University.

## Author Contributions

Conceptualization, M.H., Y.M., M.S. and H.B.L.; Methodology, M.H., Y.M., Z.L., D.C., N.L., M.S. and H.B.L.; Experimental Execution, M.H., Y.M., D.C., N.L., W.D.M., W.M. and S.J.; Software, M.H., Y.M., Z.L. K.S., L.C. L.J; Formal Analysis and interpretation, M.H., Y.M. and Z.L.; Figure-Making, M.H., Y.M. and Z.L.; Writing, M.H., and H.B.L.; Supervision, X.D., H.Z., J.J.G.,M.S. and H.B.L.; Funding Acquisition, H.B.L. and M.S. Xinzhong Dong is an investigator for the Howard Hughes Medical Institute. All authors reviewed the paper.

