## Supplemental Information for "Molecular Display of the Animal Meta-Venome for Discovery of Novel Therapeutic Peptides"

**This PDF file includes:**

Figures S1 to S5
Tables S1 to S7

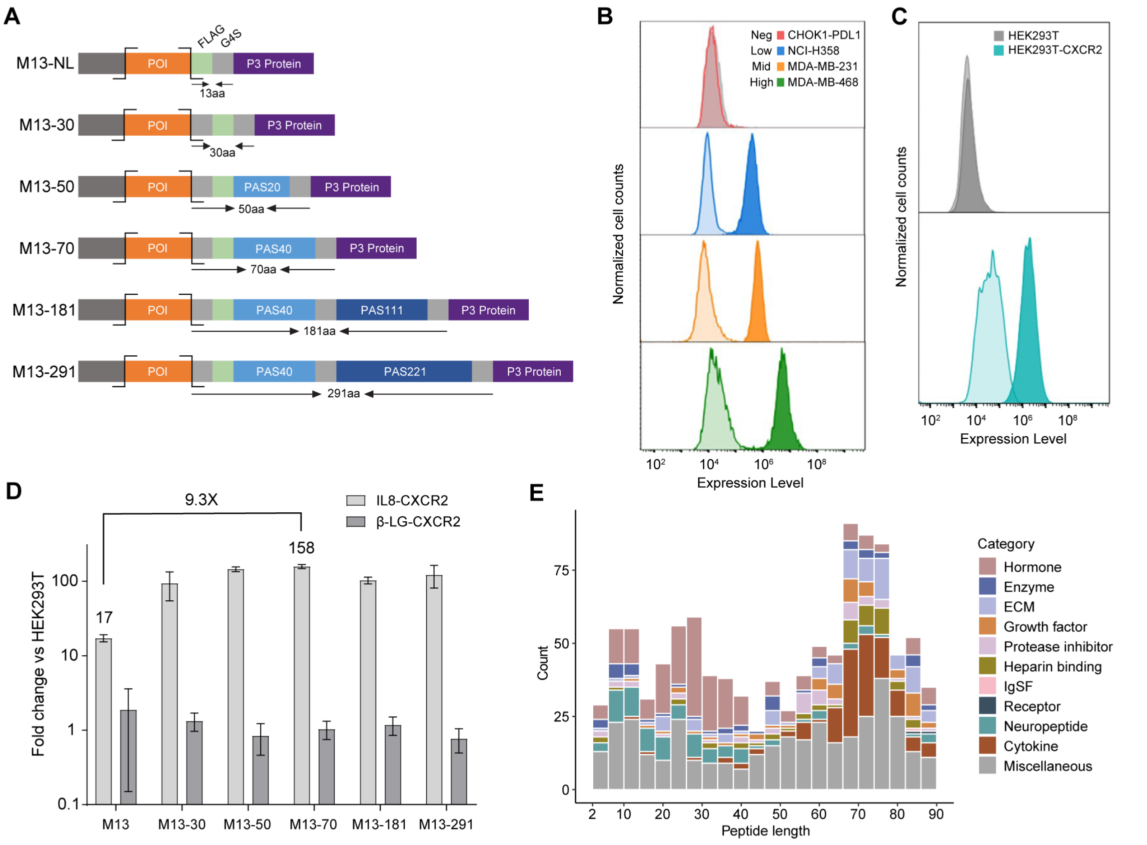


**Fig. S1. M13 hyperphage display platform development and validation.** (**A**) Depiction of the M13 phagemid vector with the key components labelled: POI, peptide of interest (orange), Flag tag (green), G4S linker (grey), PAS linkers (blue), P3 protein (purple), and restriction digestion sites (black lines). The linker lengths range from 13 (NL) to 291 amino acids. (**B**) Flow cytometry analysis of EGFR expression levels on four cell lines: CHOK1-PDL1 (negative control cell line), NCI-H358 (low), MDA-MB-231 (medium), and MDA-MB-468 (high). The lower expression level peaks represent background staining with only the fluorescent anti-mouse mAb, and the higher expression level peaks represent cells stained with anti-EGFR mAb (primary mAb) and detected with a fluorescent anti-mouse mAb. (**C**) Flow cytometry analysis of CXCR2 expression levels on HEK293T cells (CXCR2-) and HEK293T-CXCR2 (CXCR2+) cells. The lower expression level peaks represent background staining with only the fluorescent anti-mouse mAb, and the higher expression level peaks represent cells stained with anti-CXCR2 mAb (primary mAb) and detected with a fluorescent anti-mouse mAb. (**D**) Graph depicting the impact of linker length on the fold change value of IL-8 binding to CXCR2. β-Lactoglobulin (β-LG), serving as a negative control protein, exhibits low non-specific binding to HEK293T-CXCR2 cells. (**E**) Composition of the human secretome library. The histogram depicts the distribution of annotated protein molecular function versus amino acid length. The length of peptides ranges from 2 to 90 amino acids, split into 22 bins with width of 4 amino acids each bin.

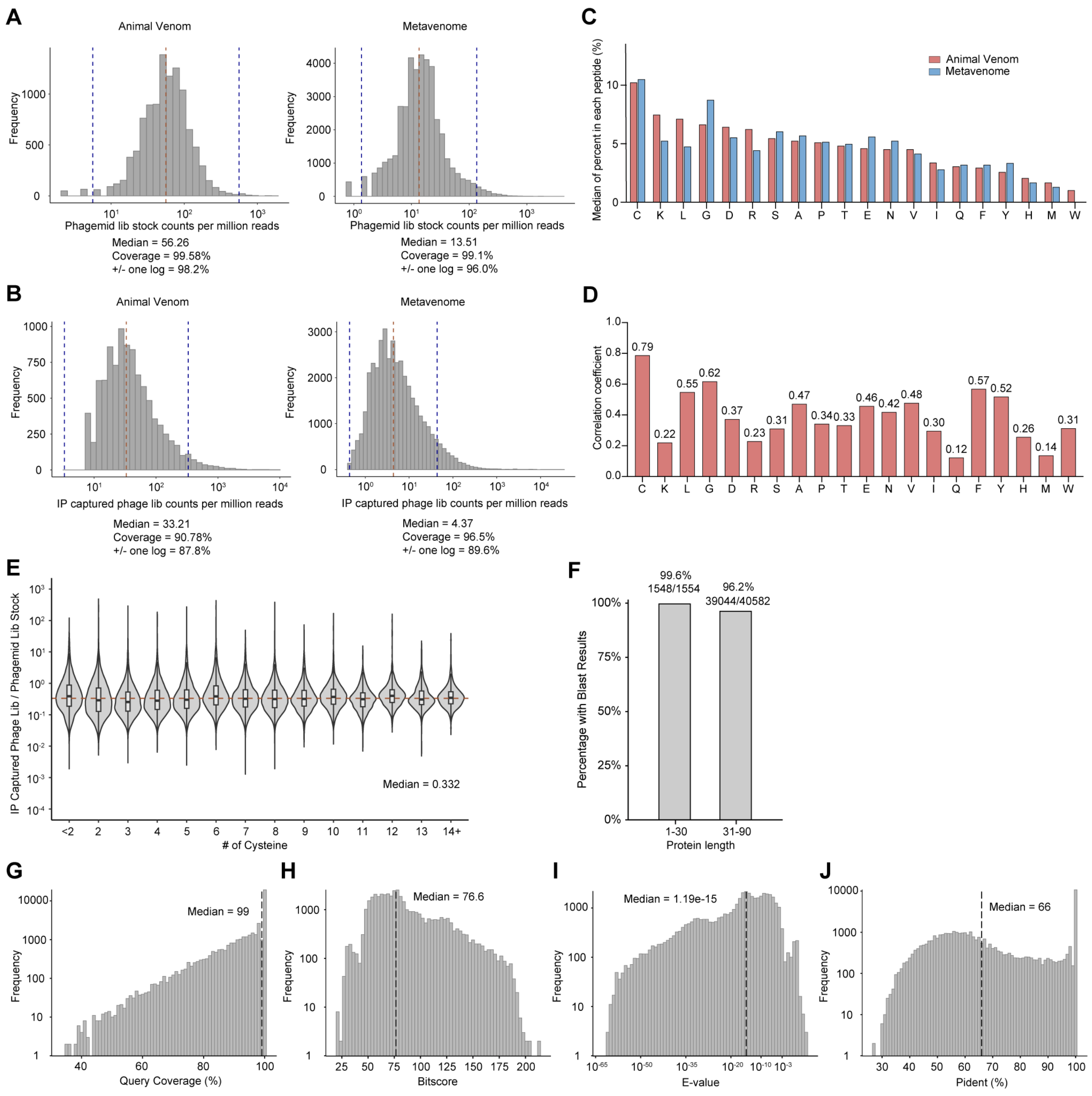


**Fig. S2. Animal venom and metavenome library quality control and characterization.** (**A**) Animal venom and Metavenome phagemid library stock quality control. For animal venom library, 99.58% of library members were detected; 98.2% of the library was within one log of the mean (indicated by vertical dashed lines). For metavenome library, 99.1% of library members were detected; 96.0% of the library was within one log of the mean. (**B**) Animal venom and Metavenome phage library quality control. For animal venom library, 90.78% of library members were detected; 87.8% of the library was within one log of the mean. For metavenome library, 96.5% of library members were detected; 89.6% of the library was within one log of the mean. (**C**) The abundance of each type of amino acid in the peptide sequences from both animal venom and metavenome libraries. Each set of bars represents a unique amino acid, with their heights corresponding to their relative abundance in the analyzed peptide sequences. (**D**) Analysis of amino acid abundance and correlation between animal venom and metavenome peptide sequence pairs. The Spearman correlation coefficient calculated from protein sequence pairs, with each bar corresponding to a distinct amino acid. The correlation coefficient is derived based on the quantity of each amino acid present in the animal venom and metavenome protein sequence pairs. (**E**) Assessment of the impact of the cysteine count on the metavenome phage library in comparison to its phagemid library stock, per number of expected cysteines. The same experimental method described for Figure 2E was used. (**F-J**) Metavenome library characterization with NCBI-BLAST+. (F) showed Library members annotated from NCBI-BLAST+. (G-J) showed essential parameters from NCBI-BLAST+ search outcomes with black dotted line in each plot highlighting the median value

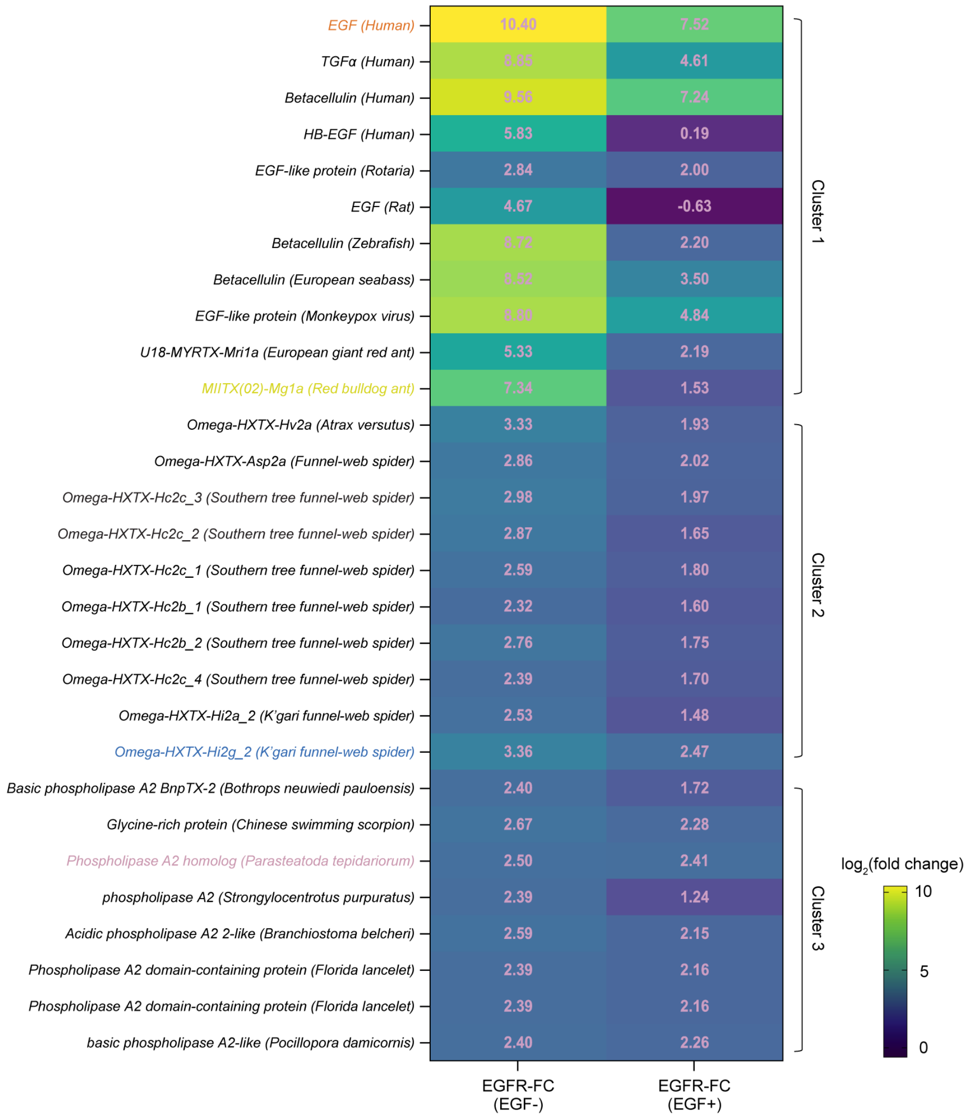


**Fig. S3. Enrichment of EGFR ligands in cluster 1-3 from the animal venom and metavenome libraries.** Fold change values for ligands binding to EGFR across clusters 1-3 are displayed in each heatmap cell. The left column presents screening results in the absence of EGF, while the right column depicts those obtained in the presence of competing EGF.

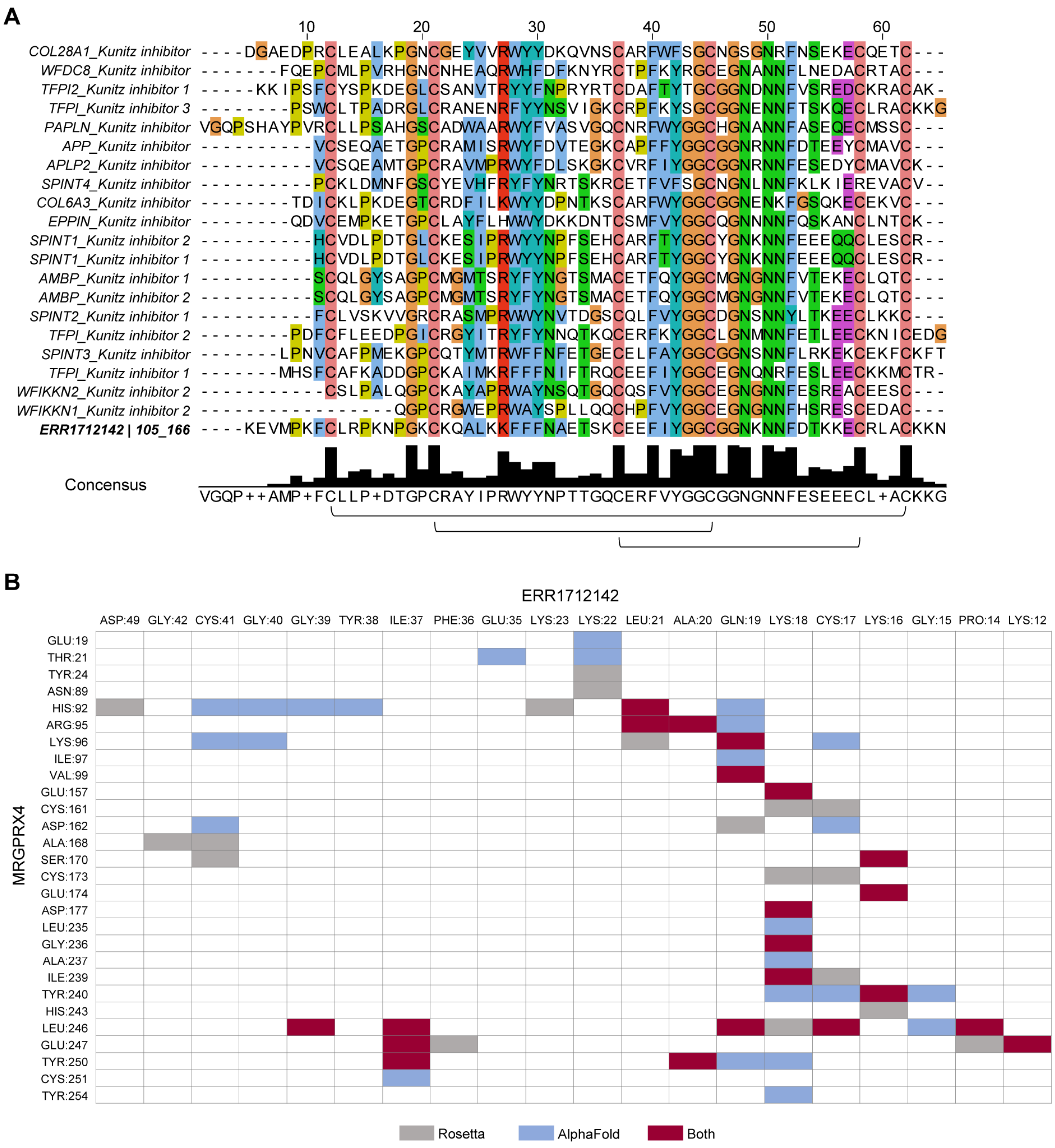


**Fig. S4. Human structural Homologs to ERR1712142|105-166 identified with Foldseek and ERR1712142|105-166 docking result.** (**A**) MSA analysis illustrates conserved amino acids shared between the 20 homologs and ERR1712142|105-166. Black underlines represent the disulfide bond patterns as annotated in the UniProt database. (**B**) Contact map of ERR1712142|105-166 -MRGPRX4 docking results via RosettaDock and AlphaFold. Amino acids between ERR1712142|105-166 and MRGPRX4 within a 5 Å radius are colored; consensus predictions between both modeling methods are indicated in red, while unique RosettaDock and AlphaFold predictions are displayed in grey and blue, respectively.

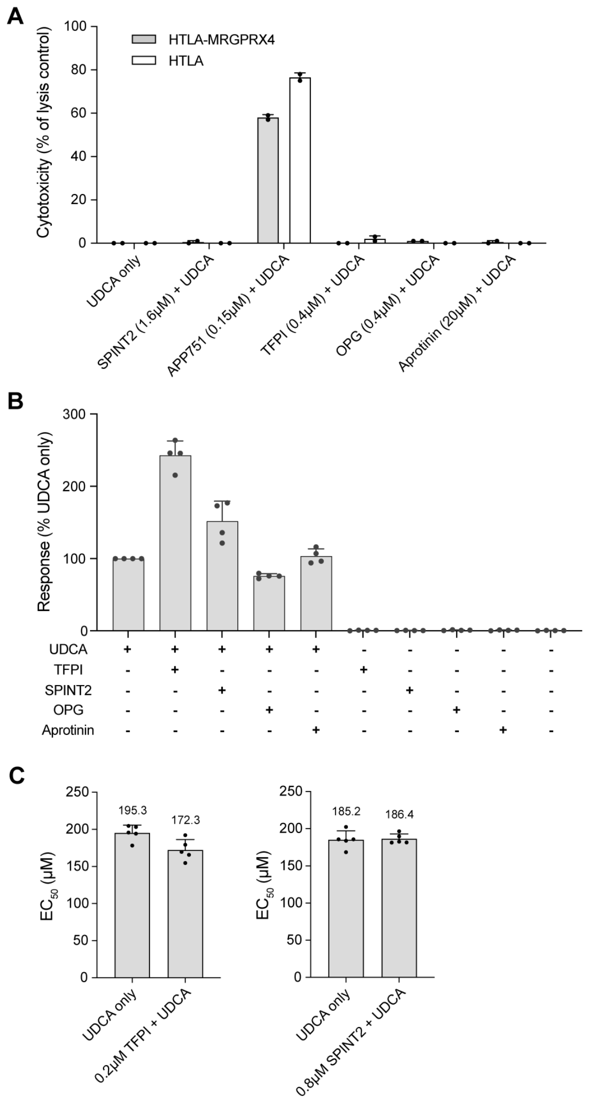


**Fig. S5. Additional results for PRESTO-Tango and cytotoxicity assays.** (**A**) Cytotoxicity of the proteins tested on PRESTO-Tango assay alongside UDCA on HTLA-MRGPRX4 and HTLA cells. Data are presented as the percentage relative to lysed cell control (mean ± SD, n=2). (**B**) Relative luminescence signal on HTLA-MRGPRX4 cells stimulated by 200μM UDCA, 0.2μM TFPI, 0.8μM SPINT2, and two negative control proteins, 0.4μM osteoprotegerin (OPG) and 20μM aprotinin on PRESTO-Tango assay. Data are presented as the percentage relative to UDCA only (mean ± SD, n=4). (**C**) $EC_{50}$ of UDCA with or without 0.2μM TFPI or 0.8μM SPINT2 (mean ± SD, n=5).

### Table S1. Sequences of tested M13 linker designs

| Linker name | Short linker | FLAG_EK | PAS linker | GS linker | Addition PAS linker | Common |
|---|---|---|---|---|---|---|
| **M13-30 (30 aa)** | GGGGS | DYKDDDDK | X | (G4S)3 | X | AS |
| **M13-50 (50aa)** | GGGGS | DYKDDDDK | **PAS20:** ASPAAPAPAS PAAPAPSAPA | (G4S)3 | X | AS |
| **M13-70 (70aa)** | GGGGS | DYKDDDDK | **PAS40:** ASPAAPAPAS PAAPAPSAPA ASPAAPAPAS PAAPAPSAPA | (G4S)3 | X | AS |
| **M13-181 (181aa)** | GGGGS | DYKDDDDK | **PAS40:** ASPAAPAPAS PAAPAPSAPA ASPAAPAPAS PAAPAPSAPA | (G4S)3 | APR - [PAS38 - (G4S)3 - PS]1 - PAS38 - (G4S)3 | AS |
| **M13-291 (291aa)** | GGGGS | DYKDDDDK | **PAS40:** ASPAAPAPAS PAAPAPSAPA ASPAAPAPAS PAAPAPSAPA | (G4S)3 | APR - [PAS38 - (G4S)3 - PS]3 - PAS38 - (G4S)3 | AS |

### Table S2. Composition of the human secretome library

| | Secretome database | | Human secretome lib | |
|---|---|---|---|---|
| | # of peptides | Proportion in original lib set (%) | # of peptides | Proportion in original lib set (%) |
| **Enzyme** | 1050 | 16.38 | 45 | 5.10 |
| **ECM** | 991 | 15.46 | 85 | 9.64 |
| **Cytokine** | 368 | 5.74 | 121 | 13.72 |
| **Hormone** | 303 | 4.73 | 200 | 22.68 |
| **Growth factor** | 286 | 4.46 | 48 | 5.44 |
| **Heparin binding** | 209 | 3.26 | 48 | 5.44 |
| **Protease inhibitor** | 199 | 3.10 | 44 | 4.99 |
| **Neuropeptide** | 112 | 1.75 | 84 | 9.52 |
| **Receptor** | 102 | 1.59 | 2 | 0.23 |
| **IgSF** | 49 | 0.76 | 1 | 0.11 |
| **Miscellaneous** | 3523 | 54.96 | 372 | 42.18 |

### Table S3. Animal venom library composition

| Phylum | Class | # of lib members |
|---|---|---|
| Mollusca (46.83%) | Gastropoda | 4962 |
| | Cephalopoda | 1 |
| Arthropoda (34.98%) | Arachnida | 3285 |
| | Insecta | 272 |
| | Chilopoda | 148 |
| | Branchiopoda | 1 |
| | Merostomata | 1 |
| Chordata (14.92%) | Lepidosauria | 1458 |
| | Amphibia | 74 |
| | Actinopteri | 20 |
| | Mammalia | 15 |
| | Aves | 12 |
| | Chondrichthyes | 2 |
| Cnidaria (3.11%) | Anthozoa | 329 |
| | Scyphozoa | 1 |
| Platyhelminthes (0.06%) | Rhabditophora | 3 |
| | Trematoda | 2 |
| | Cestoda | 1 |
| Nemertea (0.04%) | Pilidiophora | 4 |
| Annelida (0.03%) | Polychaeta | 3 |
| Porifera (0.02%) | Demospongiae | 2 |
| Nematoda (0.01%) | Enoplea | 1 |

### Table S4. Animal venom library quality analysis

| | Animal toxin lib db | Phagemid library stock | | IP captured phage lib | |
|---|---|---|---|---|---|
| # of Disulfide bonds | # of peptides | # of peptides | Coverage | # of peptides | Coverage |
| 0 | 1033 | 1019 | 98.6% | 931 | 90.1% |
| 1 | 210 | 209 | 99.5% | 193 | 91.9% |
| 2 | 435 | 434 | 99.8% | 402 | 92.4% |
| 3 | 1756 | 1754 | 99.9% | 1583 | 90.1% |
| 4 | 1137 | 1131 | 99.5% | 1013 | 89.1% |
| 5 | 243 | 242 | 99.6% | 206 | 84.8% |
| 6 | 121 | 121 | 100.0% | 95 | 78.5% |
| 7 | 13 | 13 | 100.0% | 12 | 92.3% |
| 8 | 2 | 2 | 100.0% | 1 | 50.0% |
| 10 | 1 | 1 | 100.0% | 1 | 100.0% |
| Unknown (NA) | 5646 | 5626 | 99.6% | 5183 | 91.8% |
| Total | 10597 | 10552 | 99.6% | 9620 | 90.8% |

### Table S5. Metavenome library quality analysis

| | Metavenome lib db | Phagemid library stock | | IP captured phage lib | |
|---|---|---|---|---|---|
| # of Cysteine | # of peptides | # of peptides | Coverage | # of peptides | Coverage |
| 0 | 2806 | 2754 | 98.1% | 2693 | 96.0% |
| 1 | 2472 | 2447 | 99.0% | 2379 | 96.2% |
| 2 | 2722 | 2711 | 99.6% | 2621 | 96.3% |
| 3 | 3618 | 3602 | 99.6% | 3424 | 94.6% |
| 4 | 2228 | 2200 | 98.7% | 2121 | 95.2% |
| 5 | 3440 | 3409 | 99.1% | 3306 | 96.1% |
| 6 | 9704 | 9596 | 98.9% | 9433 | 97.2% |
| 7 | 3223 | 3197 | 99.2% | 3114 | 96.6% |
| 8 | 4406 | 4378 | 99.4% | 4233 | 96.1% |
| 9 | 1690 | 1674 | 99.1% | 1634 | 96.7% |
| 10 | 2130 | 2114 | 99.2% | 2087 | 98.0% |
| 11 | 813 | 809 | 99.5% | 791 | 97.3% |
| 12 | 793 | 786 | 99.1% | 783 | 98.7% |
| 13 | 356 | 353 | 99.2% | 351 | 98.6% |
| 14 | 476 | 476 | 100.0% | 476 | 100.0% |
| 15 | 188 | 187 | 99.5% | 186 | 98.9% |
| 16 | 61 | 60 | 98.4% | 59 | 96.7% |
| 17 | 5 | 5 | 100.0% | 5 | 100.0% |
| 19 | 5 | 5 | 100.0% | 5 | 100.0% |
| Total | 41136 | 40763 | 99.1% | 39701 | 96.5% |

### Table S6. Metavenome library composition: top 10 phyla

| Phylum | Class | # of lib members |
|---|---|---|
| Arthropoda (27.43%) | Arachnida | 5826 |
| | Insecta | 3241 |
| | Hexanauplia | 877 |
| | Malacostraca | 569 |
| | Collembola | 460 |
| | Branchiopoda | 275 |
| | Ostracoda | 97 |
| | Merostomata | 76 |
| | Thecostraca | 70 |
| | Chilopoda | 42 |
| | Pycnogonida | 20 |
| | Diplopoda | 1 |
| | None | 1 |
| | Symphyla | 1 |
| Chordata (24.46%) | Actinopteri | 2653 |
| | Mammalia | 2514 |
| | Lepidosauria | 2445 |
| | Aves | 811 |
| | None | 494 |
| | Amphibia | 381 |
| | Leptocardii | 370 |
| | Appendicularia | 210 |
| | Ascidiacea | 199 |
| | Chondrichthyes | 124 |
| | Hyperoartia | 60 |
| | Cladistia | 45 |
| | Myxini | 1 |
| Mollusca (10.71%) | Gastropoda | 3108 |
| | Bivalvia | 1329 |
| | Cephalopoda | 75 |
| Proteobacteria (5.03%) | Deltaproteobacteria | 1163 |
| | Gammaproteobacteria | 370 |
| | Alphaproteobacteria | 275 |
| | Betaproteobacteria | 203 |
| | None | 70 |
| | Epsilonproteobacteria | 19 |
| | Oligoflexia | 16 |
| | Hydrogenophilalia | 3 |
| | Acidithiobacillia | 1 |
| | Candidatus Muproteobacteria | 1 |
| Cnidaria (4.74%) | Anthozoa | 1864 |
| | Hydrozoa | 109 |
| | Scyphozoa | 16 |
| | Myxozoa | 6 |
| | Cubozoa | 2 |
| Nematoda (3.76%) | Chromadorea | 1202 |
| | Enoplea | 381 |

### Table S6. (continued)

| Phylum | Class | # of lib members |
|---|---|---|
| Rotifera (2.21%) | Eurotatoria | 924 |
| | Pararotatoria | 9 |
| Platyhelminthes (1.68%) | Cestoda | 403 |
| | Trematoda | 214 |
| | Rhabditophora | 83 |
| | Monogenea | 9 |
| Ascomycota (1.68%) | Sordariomycetes | 200 |
| | Eurotiomycetes | 159 |
| | Dothideomycetes | 120 |
| | Leotiomycetes | 117 |
| | Pezizomycetes | 50 |
| | Lecanoromycetes | 31 |
| | Geoglossomycetes | 8 |
| | Saccharomycetes | 8 |
| | Orbiliomycetes | 5 |
| | Pneumocystidomycetes | 4 |
| | Neolectomycetes | 2 |
| | Schizosaccharomycetes | 1 |
| | Taphrinomycetes | 1 |
| Annelida (1.37%) | Polychaeta | 441 |
| | Clitellata | 135 |

### Table S7. Structural Homologs to ERR1712142|105-166 in Human Proteins Identified with Foldseek.

| Target ID | uniprot_ID | Seq.id. | Query Pos_start | Query Pos_end | Target Pos_start | Target Pos_end | E-Value | Score | Query seq. | Target seq. | Protein | Domain | Gene |
|---|---|---|---|---|---|---|---|---|---|---|---|---|---|
| AF-P05067-F1-model_v4 | P05067 | 40.3 | 7 | 58 | 290 | 341 | 1.71E-05 | 257 | FCLRPKNPGKCKQALKKFFFNAETSKCEEFIYGGCGGNKNNFDTKKECRLAC | VCSEQAETGPCRAMISRWYFDVTEGKCAPFFYGGCGGNRNNFDTEEYCMAVC | Amyloid-beta precursor protein | BPTI/Kunitz inhibitor | APP (A4, AD1) |
| AF-Q06481-F1-model_v4 | Q06481 | 39.6 | 7 | 59 | 309 | 361 | 1.71E-05 | 258 | FCLRPKNPGKCKQALKKFFFNAETSKCEEFIYGGCGGNKNNFDTKKECRLACK | VCSQEAMTGPCRAVMPRWYFDLSKGKCVRFIYGGCGGNRNNFESEDYCMAVCK | Amyloid-like protein 2 | BPTI/Kunitz inhibitor | APLP2 |
| AF-Q8TEU8-F1-model_v4 | Q8TEU8 | 39.2 | 8 | 58 | 386 | 436 | 3.40E-04 | 208 | CLRPKNPGKCKQALKKFFFNAETSKCEEFIYGGCGGNKNNFDTKKECRLAC | CSLPALQGPCKAYAPRWAYNSQTGQCQSFVYGGCEGNGNNFESREACEESC | WAP, Kazal, immunoglobulin, Kunitz and NTR domain-containing protein 2 | BPTI/Kunitz inhibitor 2 | WFIKKN2 |
| AF-P49223-F1-model_v4 | P49223 | 46.5 | 4 | 61 | 32 | 89 | 4.08E-06 | 286 | MPKFCLRPKNPGKCKQALKKFFFNAETSKCEEFIYGGCGGNKNNFDTKKECRLACKKN | LPNVCAFPMEKGPCQTYMTRWFFNFETGECELFAYGGCGGNSNNFLRKEKCEKFCKFT | Kunitz-type protease inhibitor 3 | BPTI/Kunitz inhibitor | SPINT3 |
| AF-P48307-F1-model_v4 | P48307 | 40.6 | 2 | 60 | 152 | 210 | 1.1E-06 | 304 | EVMPKFCLRPKNPGKCKQALKKFFFNAETSKCEEFIYGGCGGNKNNFDTKKECRLACKK | KKIPSFCYSPKDEGLCSANVTRYYFNPRYRTCDAFTYTGCGGNDNNFVSREDCKRACAK | Tissue factor pathway inhibitor 2 | BPTI/Kunitz inhibitor 1 | TFPI2 |
| AF-O43291-F1-model_v4 | O43291 | 42.3 | 7 | 58 | 37 | 88 | 2.97E-05 | 269 | FCLRPKNPGKCKQALKKFFFNAETSKCEEFIYGGCGGNKNNFDTKKECRLAC | FCLVSKVVGRCRASMPRWWYNVTDGSCQLFVYGGCDGNSNNYLTKEECLKKC | Kunitz-type protease inhibitor 2 | BPTI/Kunitz inhibitor 1 | SPINT2 |
| AF-P10646-F1-model_v4 | P10646 | 49.1 | 4 | 60 | 50 | 106 | 1.60E-06 | 301 | MPKFCLRPKNPGKCKQALKKFFFNAETSKCEEFIYGGCGGNKNNFDTKKECRLACKK | MHSFCAFKADDGPCKAIMKRFFFNIFTRQCEEFIYGGCEGNQNRFESLEECKKMCTR | Tissue factor pathway inhibitor | BPTI/Kunitz inhibitor 1 | TFPI |
| 1tfx_D | P10646 | 43.8 | 5 | 61 | 2 | 58 | 1.61E-06 | 283 | PKFCLRPKNPGKCKQALKKFFFNAETSKCEEFIYGGCGGNKNNFDTKKECRLACKKN | PDFCFLEEDPGICRGYITRYFYNNQTKQCERFKYGGCLGNMNNFETLEECKNICEDG | Tissue factor pathway inhibitor | BPTI/Kunitz inhibitor 2 | TFPI |
| 1irh_A | P10646 | 47.3 | 5 | 61 | 5 | 61 | 2.22E-04 | 187 | PKFCLRPKNPGKCKQALKKFFFNAETSKCEEFIYGGCGGNKNNFDTKKECRLACKKN | PSWCLTPADRGLCRANENRFYYNSVIGKCRPFKYSGCGGNENNFTSKQECLRACKKG | Tissue factor pathway inhibitor | BPTI/Kunitz inhibitor 3 | TFPI |
| AF-P02760-F1-model_v4 | P02760 | 42.3 | 7 | 58 | 230 | 281 | 3.39E-05 | 246 | FCLRPKNPGKCKQALKKFFFNAETSKCEEFIYGGCGGNKNNFDTKKECRLAC | SCQLGYSAGPCMGMTSRYFYNGTSMACETFQYGGCMGNGNNFVTEKECLQTC | Protein AMBP | BPTI/Kunitz inhibitor 1 | AMBP |
| AF-O95925-F1-model_v4 | O95925 | 36.3 | 5 | 59 | 74 | 128 | 8.64E-05 | 231 | PKFCLRPKNPGKCKQALKKFFFNAETSKCEEFIYGGCGGNKNNFDTKKECRLACK | QDVCEMPKETGPCLAYFLHWWYDKKDNTCSMFVYGGCQGNNNNFQSKANCLNTCK | Eppin | BPTI/Kunitz inhibitor | EPPIN |
| AF-Q96NZ8-F1-model_v4 | Q96NZ8 | 35.5 | 14 | 58 | 365 | 409 | 0.003016 | 166 | PGKCKQALKKFFFNAETSKCEEFIYGGCGGNKNNFDTKKECRLAC | QGPCRGWEPRWAYSPLLQQCHPFVYGGCEGNGNNFHSRESCEDAC | WAP, Kazal, immunoglobulin, Kunitz and NTR domain-containing protein 1 | BPTI/Kunitz inhibitor 2 | WFIKKN1 |

### Table S7. (continued)

| Target ID | uniprot_ID | Seq.id. | Query Pos_start | Query Pos_end | Target Pos_start | Target Pos_end | E-Value | Score | Query seq. | Target seq. | Protein | Domain | Gene |
|---|---|---|---|---|---|---|---|---|---|---|---|---|---|
| AF-Q8IUA0-F1-model_v4 | Q8IUA0 | 34.5 | 4 | 58 | 91 | 145 | 4.09E-05 | 232 | MPKFCLRPKNPGKCKQALKKFFFNAETSKCEEFIYGGCGGNKNNFDTKKECRLAC | FQEPCMLPVRHGNCNHEAQRWHFDFKNYRCTPFKYRGCEGNANNFLNEDACRTAC | WAP four-disulfide core domain protein 8 | BPTI/Kunitz inhibitor | WFDC8 |
| AF-Q2UY09-F1-model_v4 | Q2UY09 | 27.5 | 1 | 58 | 1065 | 1122 | 3.19E-05 | 229 | KEVMPKFCLRPKNPGKCKQALKKFFFNAETSKCEEFIYGGCGGNKNNFDTKKECRLAC | DGAEDPRCLEALKPGNCGEYVVRWYYDKQVNSCARFWFSGCNGSGNRFNSEKECQETC | Collagen alpha-1(XXVIII) chain | BPTI/Kunitz inhibitor | COL28A1 |
| AF-O43278-F1-model_v4 | O43278 | 35.8 | 7 | 59 | 390 | 442 | 9.79E-05 | 231 | FCLRPKNPGKCKQALKKFFFNAETSKCEEFIYGGCGGNKNNFDTKKECRLACK | HCVDLPDTGLCKESIPRWYYNPFSEHCARFTYGGCYGNKNNFEEEQQCLESCR | Kunitz-type protease inhibitor 1 | BPTI/Kunitz inhibitor 2 | SPINT1 |
| AF-Q6UDR6-F1-model_v4 Kun | Q6UDR6 | 37.7 | 7 | 59 | 40 | 92 | 1.33E-04 | 238 | FCLRPKNPGKCKQALKKFFFNAETSKCEEFIYGGCGGNKNNFDTKKECRLACK | PCKLDMNFGSCYEVHFRYFYNRTSKRCETFVFSGCNGNLNNFKLKIEREVACV | Kunitz-type protease inhibitor 4 | BPTI/Kunitz inhibitor | SPINT4 |
| AF-O95428-F1-model_v4 | O95428 | 35.4 | 2 | 58 | 743 | 804 | 1.53E-02 | 134 | EVMPKF-----CLRPKNPGKCKQALKKFFFNAETSKCEEFIYGGCGGNKNNFDTKKECRLAC | VGQPSHAYPVRCLLPSAHGSCADWAARWYFVASVGQCNRFWYGGCHGNANNFASEQECMSSC | Papilin | BPTI/Kunitz inhibitor | PAPLN |
| 1knt_A | P12111 | 40.7 | 5 | 58 | 1 | 54 | 7.69E-05 | 221 | PKFCLRPKNPGKCKQALKKFFFNAETSKCEEFIYGGCGGNKNNFDTKKECRLAC | TDICKLPKDEGTCRDFILKWYYDPNTKSCARFWYGGCGGNENKFGSQKECEKVC | Collagen alpha-3(VI) chain | BPTI/Kunitz inhibitor | COL6A3 |
| AF-P02760-F1-model_v4 | P02760 | 42.3 | 7 | 58 | 230 | 281 | 3.39E-05 | 246 | FCLRPKNPGKCKQALKKFFFNAETSKCEEFIYGGCGGNKNNFDTKKECRLAC | SCQLGYSAGPCMGMTSRYFYNGTSMACETFQYGGCMGNGNNFVTEKECLQTC | Protein AMBP | BPTI/Kunitz inhibitor 2 | AMBP |
| AF-O43278-F1-model_v4 | O43278 | 35.8 | 7 | 59 | 390 | 442 | 9.79E-05 | 231 | FCLRPKNPGKCKQALKKFFFNAETSKCEEFIYGGCGGNKNNFDTKKECRLACK | HCVDLPDTGLCKESIPRWYYNPFSEHCARFTYGGCYGNKNNFEEEQQCLESCR | Kunitz-type protease inhibitor 1 | BPTI/Kunitz inhibitor 1 | SPINT1 |
